# Evolutionary analysis reveals the origin of sodium coupling in glutamate transporters

**DOI:** 10.1101/2023.12.03.569786

**Authors:** Krishna D. Reddy, Burha Rasool, Farideh Badichi Akher, Nemanja Kutlešić, Swati Pant, Olga Boudker

## Abstract

Secondary active membrane transporters harness the energy of ion gradients to concentrate their substrates. Homologous transporters evolved to couple transport to different ions in response to changing environments and needs. The bases of such diversification, and thus principles of ion coupling, are unexplored. Employing phylogenetics and ancestral protein reconstruction, we investigated sodium-coupled transport in prokaryotic glutamate transporters, a mechanism ubiquitous across life domains and critical to neurotransmitter recycling in humans. We found that the evolutionary transition from sodium-dependent to independent substrate binding to the transporter preceded changes in the coupling mechanism. Structural and functional experiments suggest that the transition entailed allosteric mutations, making sodium binding dispensable without affecting ion-binding sites. Allosteric tuning of transporters’ energy landscapes might be a widespread route of their functional diversification.

## Introduction

Transport of solutes across cell membranes against their concentration gradients is critical for cellular homeostasis. Secondary active membrane transporters catalyze this energetically unfavorable process using the energy of ions flowing down their electrochemical gradients. The ion-binding sites of many transporters are well-characterized. However, the three-dimensional structural properties of transporters necessary to couple substrate and ion movements and thus harvest the electrochemical energy remain opaque.

In the dicarboxylate/amino acid:cation symporter family, the Na^+^-coupling mechanism persisted through millions of years, conserved between archaea and humans, where excitatory amino acid transporters (EAAT or SLC1) use Na^+^ gradients to clear the neurotransmitter glutamate from the synaptic cleft. Many transporter families, including glutamate transporters, show functional diversification using Na^+^ or H^+^ gradients (**Figure 1a**) (*1, 2*). The cell environment and needs provide the selective pressure to use specific ions, such that when cell membranes are H^+^-leaky or Na^+^ is abundant, Na^+^-coupled transport predominates. In contrast, when Na^+^ is scarce, Na^+^ coupling is no longer beneficial and is eliminated. We used phylogenetic analysis, ancestral protein reconstruction (APR), and structural and functional characterization of inferred ancestral proteins to understand how prokaryotic glutamate transporters transitioned away from Na^+^-coupling. Contrary to our expectations, the transition did not involve a promiscuous ion coupling (*3, 4*) or a gradual loss of Na^+^-binding sites. Instead, allosteric changes first made Na^+^-binding dispensable to function in a distinct evolutionary step, and only then the transporters lost their Na^+^-binding residues.

**Figure 1:**
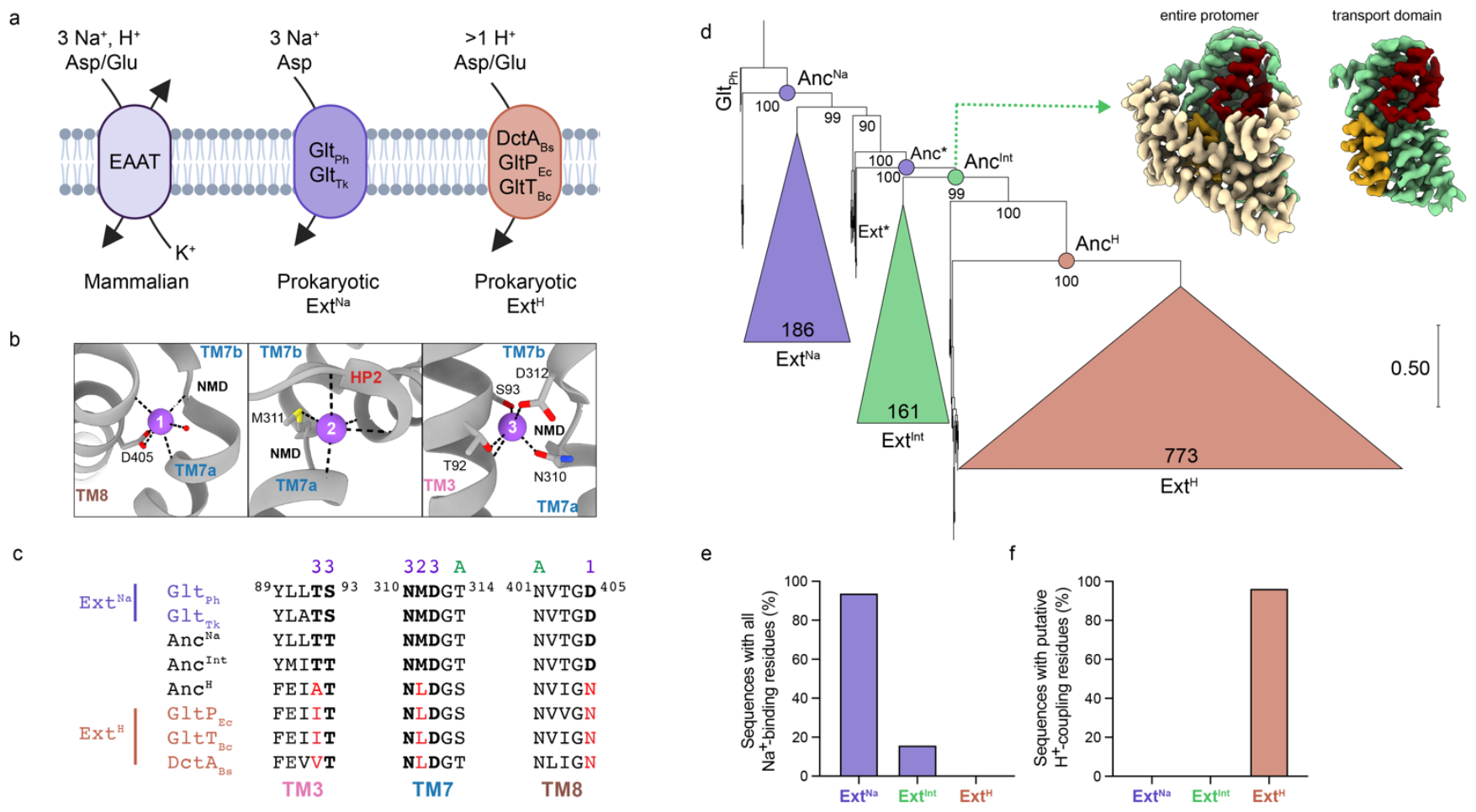
Mapping the evolutionary transition from sodium- to proton-coupling. (a) Ion-coupling stoichiometries of characterized concentrative transporters in the glutamate transporter family. (b) The architecture of Na^+^-binding sites in Glt_Ph_ (PDB 6X15). Bound Na1-3 are shown as purple spheres. Dotted lines emphasize the coordinating interactions with sidechains, shown as sticks, and main chain carbonyl oxygens, shown as cartoons. (c) Sequence alignment of experimentally characterized prokaryotic homologs and inferred ancestral transporters from Figure 1d. The numbers above the alignment refer to the Na^+^ binding sites, and ‘A’ refers to sidechains coordinating the substrate L-Asp. Signature sequence changes associated with the transition to H^+^-coupling are shown in red. (d) Reduced ML phylogeny of glutamate transporters in *Firmicutes*. The collapsed Ext^Na^ (purple), Ext^Int^ (green), and Ext^H^ (orange) clades are proportional to the number of the comprising sequences, which is shown within the clade. Numbers at nodes represent corresponding UFBoot2 values. Nodes corresponding to the reconstructed ancestral sequences are highlighted in the corresponding color and labeled Anc^Na^, Anc*, Anc^Int^, and Anc^H^. Inset: representative cryo-EM structure of an Anc^Int^ protomer in lipid nanodiscs. Scaffold domain in wheat, HP1 in yellow, HP2 in red. (e) Percentage of sequences in each clade containing all Na^+^-binding site residues ([S/T]92-93, N310, [M/S]311, D312, D405). (f) Percentage of sequences in each clade containing signature H^+^-coupling residues ([A/L/I/V/M]92, L311, N405).

Energetically linked substrate and Na^+^ binding underlies symport in Na^+^-coupled glutamate transporter homologs, including archaeal Glt_Ph_ and Glt_Tk_. A higher extracellular Na^+^ concentration ensures a greater probability of substrate binding from the outside and release on the inside of the membrane, providing directionality to the transport cycle and leading to concentrative uptake. The binding sites for the substrate and three symported Na^+^ ions are located within a dynamic transport domain, which moves between extracellular and intracellular positions in an elevator-like motion (outward- and inward-facing states, **Supplementary Figure 1**) relative to a stationary scaffold/trimerization domain (*13–22*). The transport mechanism and homotrimeric architecture are conserved across glutamate transporters (*23–27*). Na^+^ binding to two sites (Na1 and Na3) induces the formation of a high-affinity substrate binding pocket, resulting in substrate and the third Na^+^ (Na2) binding and closure of the helical hairpin 2 (HP2) gate, which precedes translocation (*7, 11, 28–32*) (**Figure 1b-c**). Thus, coupled binding and transport occur because the high-affinity substrate binding state is a high-energy state only accessible when Na^+^ binds. Accordingly, mutations in all three Na^+^-binding sites diminish coupled substrate binding and transport (*3, 6, 33–36*).

Using APR, we reconstructed and characterized ancestral transporter sequences (**Figure 1d**); we found that an intermediate ancestral transporter (Anc^Int^) between Na^+^-coupled and H^+^-coupled ancestral transporters showed Na^+^-independent substrate binding, despite preserved Na^+^-binding sites. Cryo-EM structures showed that Anc^Int^ accessed the high-affinity substrate-binding state spontaneously, suggesting that its energy is lower than in Na^+^-coupled transporters and explaining why Na^+^ binding was no longer necessary. Detailed evolutionary analysis revealed that the shifted energetic balance from the low-to the high-affinity states was due to allosteric changes distant from the ion- and substrate-binding sites and mutations of just two residues in Anc^Int^ restored Na^+^-coupling. We propose that allosterically changing the relative energy of functional states in transporters constitutes an evolutionary mechanism allowing the diversification of strictly specific ion coupling observed in present-day transporters. Our work reveals key principles underlying Na^+^-coupling, showcases the feasibility and power of the structure-function APR in untangling the complex mechanisms of membrane transporters, and suggests a general evolutionary mechanism underlying adaptations to different ionic environments in secondary active transporters.

## Results

### An evolutionary intermediate between Na^+^- and H^+^-coupled transporters

We used phylogenetic analysis and APR to understand how bacterial glutamate transporters might have changed their ion-coupling preference (**Supplementary Figure 2**). The approach can reveal minimal subsets of stepwise, chronologically ordered changes required to alter functions, including epistatic changes, which are insufficient for the functional switch but are prerequisites for it (*37*). Subsequent inference of ancestral sequences along an evolutionary pathway provides a platform to test evolutionary and mechanistic hypotheses (*38–44*). We selected sequences in *Firmicutes/Bacillota* phylum to recapitulate the transition from Na^+^ to H^+^ coupling, rooting the phylogenetic tree to archaeal Na^+^-coupled Glt_Ph_ (Methods, **Supplementary Figure 3a-c**). The tree clustered into three major clades (**Figure 1d**) with two, predictably, displaying conserved signature sequences of Na^+^ and H^+^ coupling (**Figure 1e-f, Supplementary Figure 3d**). We refer to these present-day extant sequences as Ext^Na^ and Ext^H^ clades. Strikingly, we also observed an intermediate clade Ext^Int^ between Ext^Na^ and Ext^H^ clades. Only 16% of its sequences contain all Na^+^-coordinating residues, and none contain all H^+^-coupling signature residues (**Figure 1e-f, Supplementary Figure 3d**). This branch was robust to perturbation and reconstructed with high confidence based on UFBoot2 bootstrap value > 95% (**Figure 1d**) (*45*).

**Figure 2:**
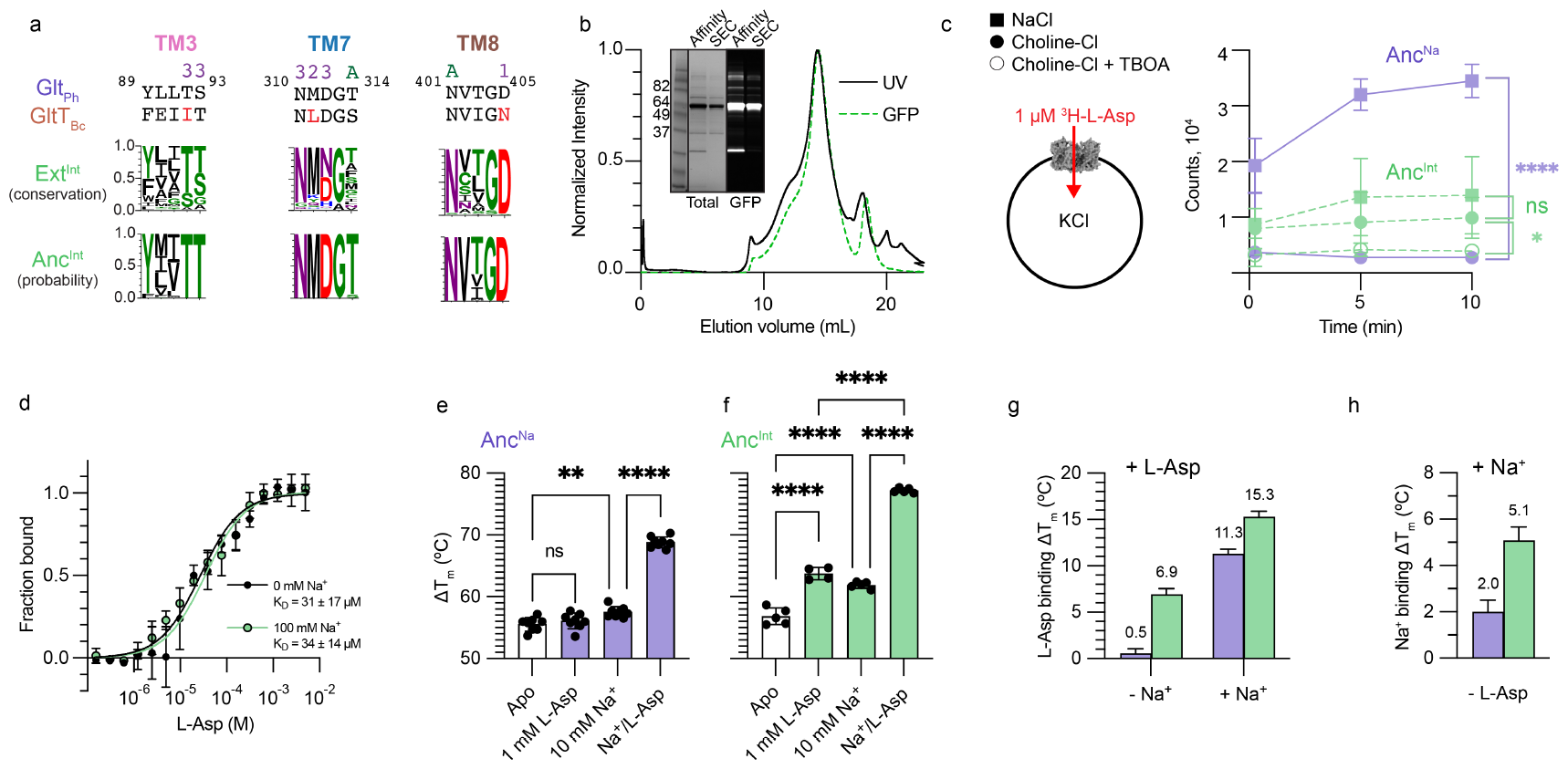
Characterization of an Na^+^-independent ancestral transporter. (a) Comparison of the signature Na^+^-coupling residues in present-day Ext^Int^ transporters and inferred Anc^Int^. Ext^Int^ WebLogo represents the conservation of transporter residues in the Ext^Int^ clade. Anc^Int^ WebLogo is a visual representation of the posterior probability per site in the ancestral reconstruction; the topmost residue at each site represents the ‘maximum-likelihood’ residue. (b) Representative size-exclusion chromatography profiles of GFP-tagged Anc^Int^ purified in apo conditions (50 mM KH_2_PO_4_/K_2_HPO_4_, 1 mM DDM). Black and dotted green lines correspond to relative optical absorption at 280 nm and GFP fluorescence excited at 485nm and measured at 510 nm, respectively. Inset: SDS PAGE analysis of GFP-tagged Anc^Int,^ stained with total protein Coomassie stain (Total) or imaged by in-gel GFP fluorescence (GFP). “Affinity” refers to pooled and concentrated elution fractions following streptactin affinity purification. “SEC” refers to pooled and concentrated peak fractions eluting at 14.5 mL following size-exclusion chromatography. (c) ^3^H-L-Asp uptake in the presence (filled squares) or absence (filled circles) of a Na^+^ gradient. A simplified proteoliposome reaction setup is on the left. All experiments were performed in the presence of a negative membrane voltage generated by valinomycin-mediated K^+^ efflux. Lines represent Anc^Na^ (solid purple) or Anc^Int^ (dashed green). Open circles represent Anc^Int^ without a Na^+^ gradient and in the presence of 1 mM DL-TBOA. Statistical significance was measured with unpaired t-tests at 10 minutes. Results were averaged from three independent experiments. (d) L-Asp binding by Anc^Int^ in 100 mM NaCl (green circles) or 100 mM choline chloride (black circles), measured via microscale thermophoresis. *K*_*D*_ in NaCl and choline chloride were 31 ± 17 μM and 34 ± 14 μM, respectively. Results are from three independent experiments. (e-f) Melting temperatures (*T*_*m*_) of Anc^Na^ (e) or Anc^Int^ (f) in buffer alone (Apo), 10 mM NaCl, 1 mM L-Asp, or both NaCl and L-Asp. Replicates are from at least two independent experiments; statistical significance was measured with ordinary one-way ANOVAs and Tukey’s multiple comparisons test. (g) Δ*T*_*m*_ of L-Asp for Anc^Na^ (purple) or Anc^Int^ (green) in the absence (-Na^+^) or presence (+ Na^+^) of sodium. Error bars are SE of the mean difference. (h) Δ*T*_*m*_ of Na^+^ for Anc^Na^ (purple) or Anc^Int^ (green) in the absence of L-Asp. Error bars are SE of the mean difference.

**Figure 3:**
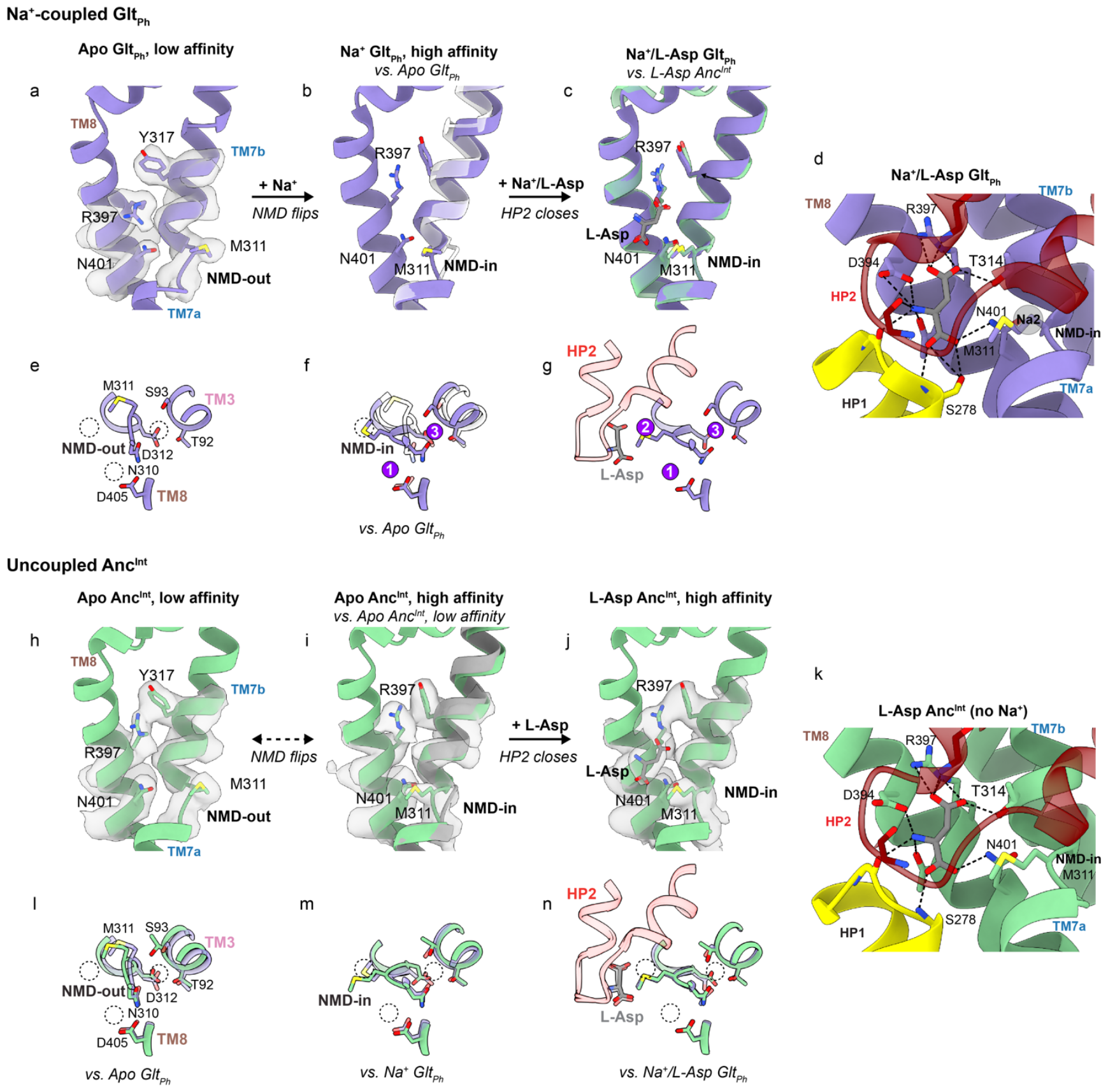
Substrate binding mechanism in Na^+^-coupled and uncoupled transporters. All numbered residues use Glt_Ph_ numbering. All structures are aligned on the HP1/TM7a region (residues 259-309 in Glt_Ph_). All densities are in grey and are sharpened and contoured at 10s. Purple spheres represent bound Na^+^ ions, and dashed circles represent unoccupied Na^+^ sites. Na^+^-coupled substrate binding of Glt_Ph_ (purple, a-g) and uncoupled substrate binding of Anc^Int^ (green, h-n) Panels depict substrate (a-d, h-k) and ion (e-g, l-n) binding sites. (a,e) In apo Glt_Ph_ (outward-facing), both substrate and ion binding sites are distorted, forming a low-affinity configuration. (b,f) Superposition of Na^+^-bound Glt_Ph_ (PDB 7AHK) and translucent apo Glt_Ph_. Upon Na1 and Na3 binding, the NMD loop (containing N310, M311, and D312) and M311 side chain flip into the pocket. TM3 (containing S92, T93) moves closer to the NMD loop to form the Na3 and Na1 sites, and M311/R397/N401 side chains adjust to form Na2 and L-Asp binding sites. These changes produce the high-affinity substrate binding state. (c-d,g) Na^+^/L-Asp bound Glt_Ph_ (PDB 6X15) demonstrates subsequent L-Asp/Na2 binding and HP2 closure; L-Asp coordination determined by hbonds in ChimeraX. (h-i,l-m) In apo conditions, Anc^Int^ visits both low-affinity (h,l) and high-affinity (i,m) states. Spontaneous NMD loop and side-chain movements are similar to those induced by Na^+^ binding in Glt_Ph_ (translucent purple, l-m). (j-k,n) L-Asp binding to the preformed site and HP2 closure in Anc^Int^ without Na^+^; L-Asp coordination is similar to Glt_Ph_.

Next, we inferred ancestral sequences by determining posterior probabilities (PP) of each amino acid occurring at each position (site) in the multiple sequence alignment. Each node’s final maximum likelihood amino acid (ML) sequence is constructed using the most probable amino acid at every site. We reconstructed sequences corresponding to the closest ancestor to the Na^+^-coupled root (Anc^Na^) and the ancestor of the intermediate Ext^Int^ clade (Anc^Int^); the average PP per site were 0.87 and 0.85, respectively (**Supplementary Figure 4a**), which is sufficiently high for APR studies (*46, 47*). As expected, reconstructed Anc^Na^ sequences showed signature sequences of Na^+^-coupled transporters (**Figure 1c, Supplementary Figure 4b, Supplementary Figure 5**). Surprisingly, Anc^Int^ also featured the full complement of Na^+^-binding residues with high posterior probabilities, contrasting with their poor conservation among the descendent present-day transporters (**Figure 1c, Figure 2a, Supplementary Figure 4a, Supplementary Figure 5**). These observations suggest a lack of selective pressure on Na^+^-binding residues in Anc^Int^, raising the question of its functional properties.

**Figure 4.**
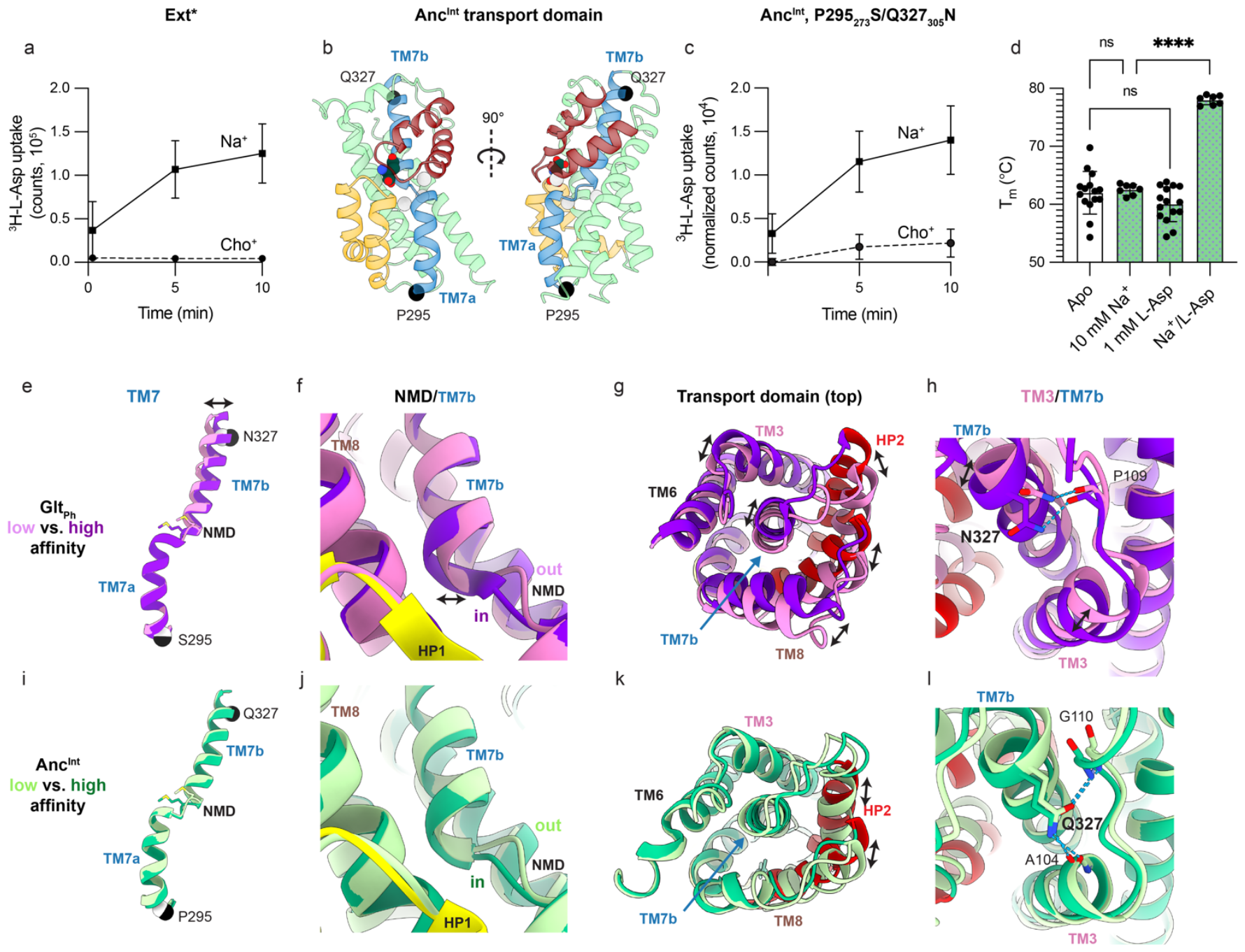
TM7 is the allosteric determinant of Na^+^ coupling. (a) ^3^H-L-Asp uptake of Ext* in the presence (filled squares) or absence (filled circles) of a Na^+^ gradient, with the same reaction setup as in Figure 2c. The data are from two independent experiments. (b) Transport domain of L-Asp bound Anc^Int^, with unoccupied Na^+^-binding sites shown as light grey spheres, TM7 in blue, and P295_273_/Q305_327_ as black spheres. (c) ^3^H-L-Asp uptake of Anc^Int^ P295_273_S/Q305_327_N with the same reaction setup as Figures 2c/4a, from three independent experiments. In each individual experiment, the first time point in choline chloride conditions (15 seconds) was set to zero. (d) Melting temperatures (*T*_*m*_) of Anc^Int^ P295_273_S/Q305_327_N in buffer alone (Apo), 10 mM NaCl, 1 mM L-Asp, or both NaCl and L-Asp. Replicates are from at least two independent experiments; statistical significance was measured with one-way ANOVAs. (e) Comparison between the low- and high-affinity Glt_Ph_ (e-h) or Anc^Int^ (i-l) structures aligned on the rigid HP1 (Glt_Ph_ residues 258-292). Darker structures are the high-affinity states (Na^+^/Asp-bound Glt_Ph_ PDB 7RCP and Asp-bound Anc^Int^), with the NMD loop positioned towards the substrate binding site. Lighter structures are apo Glt_Ph_ and low-affinity apo Anc^Int^, with the NMD loop positioned away from the substrate binding site. All residue numbering corresponds to Glt_Ph_ numbering. (e,i) TM7 alone with P295_273_/Q305_327_ shown as white and black spheres in low- and high-affinity states, respectively; view corresponds to the rotated (right) view in Figure 4b. (f,j) a close-up view of the NMD loop and the start of TM7b. Note the larger shift of TM7b in Glt_Ph_ than Anc^Int^. (g,k) a view of the transport domain from the extracellular space showing larger packing differences between low-affinity and high-affinity conformations in Glt_Ph_. (h,l) closeup view of [N/Q]327_305_ in TM7 interacting with the top of TM3 in Anc^Int^.

**Figure 5.**
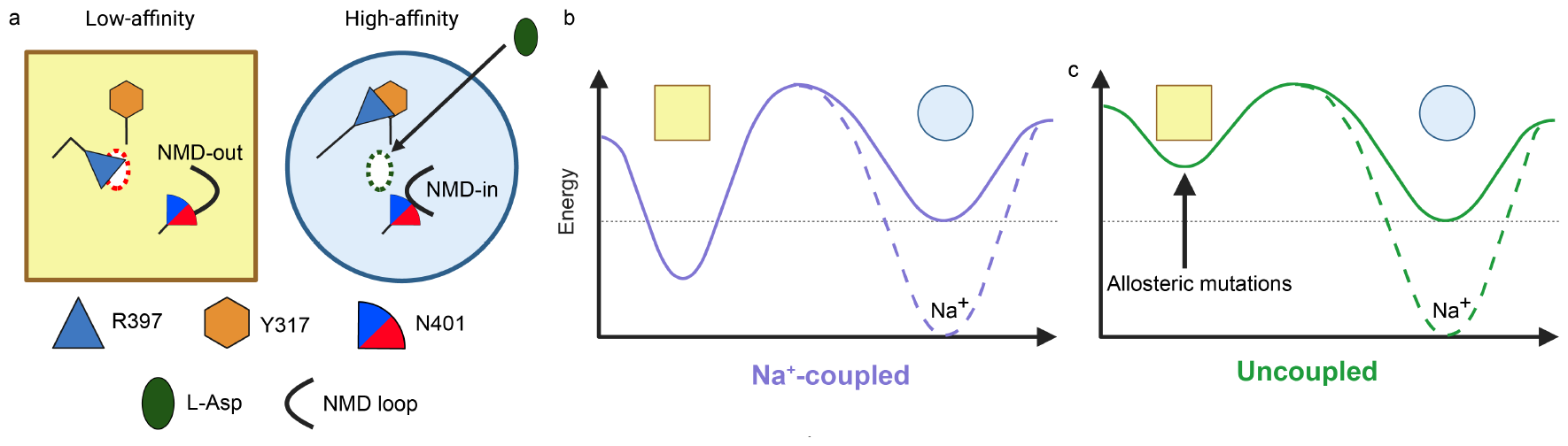
Evolution of the transporter energy landscape controls Na^+^-coupling. (a) The low-affinity (yellow square) and high-affinity (blue circle) binding configurations highlight changes in R397, N401, and the NMD loop conformations within the binding site. The square and circle shapes represent the differences in the transport domains outside the binding sites allosterically coupled to the conformations of R397, N401, and the NMD loop. The dotted ovals represent local configurations that are unsuitable (red) or ready (green) to coordinate L-Asp (green solid oval), respectively. (b-c) The hypothetical transporter energy landscapes modulated by Na^+^ binding of Na^+^-coupled (b) and uncoupled (c) transporters. Solid lines represent the relative energies of the low- and high-affinity states in the absence of Na^+^; the dashed line represents the changes upon Na^+^ binding. Relative to the high-affinity state (blue circle), the energy of the low-affinity state (yellow square) in Na^+^-coupled transporters is lower than in uncoupled transporters due to allosteric mutations outside the binding sites. In Na^+^-coupled transporters, Na^+^ binding brings the energy of the high-affinity state down, promoting substrate binding. In Na^+^-uncoupled transporters, Na^+^ binding is unnecessary.

To design ancestral constructs for protein expression, we replaced poorly aligned highly variable loops and tails, excluded from the APR analysis, with those from an extant transporter with the highest sequence identity to Anc^Int^ (**Supplementary Figure 5**). Most of these were single residue insertions, except for the C-terminus and the loop between TMs 3 and 4. These flexible regions are unlikely to affect coupling. Indeed, the C-terminus has been deleted in Glt_Ph_ for structural and functional studies (*13*), and cleavage of the loop affected transport kinetics but not ion-coupling (*48*). For consistency, we used the same loops and tails for all subsequent ancestral constructs. We expressed ancestors in *E. coli* with or without a C-terminal optimized GFP (*49*) and purified them to near homogeneity (**Figure 2b**). Despite extensive optimization, the ancestral proteins expressed ∼10-fold lower than extant prokaryotic homologs, yielding ∼20-100 μg of purified proteins per liter of culture.

### *The ancestral intermediate transporter is uncoupled from* Na^+^

When we purified and reconstituted Anc^Na^ into proteoliposomes, we observed a time-dependent accumulation of ^3^H-L-Asp in vesicles in the presence of an inward Na^+^-gradient. In contrast, no ^3^H-L-Asp was retained in the absence of Na^+^ (**Figure 2c**). These results suggest that Anc^Na^ is adequately folded and recapitulates expected Na^+^-coupling. When we repeated the experiment with Anc^Int^, we observed lower ^3^H-L-Asp retention unaffected by the removal of Na^+^ but reduced in the presence of the competitive inhibitor DL-TBOA (**Figure 2c**). We did not observe ^3^H-L-Asp retention in Anc^Int^ proteoliposomes under exchange conditions, with 500 μM unlabeled L-Asp inside the vesicles and 1 μM ^3^H-L-Asp outside (not shown). Thus, we think Anc^Int^ in proteoliposomes can bind L-Asp independently of Na^+^ but not transport it.

We used microscale thermophoresis (MST) to measure substrate binding affinity to Anc^Int^. Anc^Int^ binds L-Asp with similar micromolar affinities in the absence and presence of 100 mM Na^+^ (**Figure 2d**). Notably, the MST signal-to-noise values upon L-Asp binding ranged from 8.6 to 19.5, reflecting the high confidence of the binding isotherms (**Supplementary Figure 6a**). However, the MST signal-to-noise was insufficient to accurately determine Anc^Na^ binding affinities, likely due to changes in measurable parameters, including charge, hydration shell, or conformation (*50*). Instead, we used nano-differential scanning fluorimetry (nanoDSF) to measure ligand-dependent protein melting temperatures (*T*_*m*_) as a qualitative proxy for ligand binding (*51, 52*). In Anc^Na^, we saw insignificant or small *T*_*m*_ increases when adding 1 mM L-Asp or 10 mM Na^+^ separately compared to the apo transporter (**Figure 2e,g-h, Supplementary Figure 6b**). In contrast, we observed a dramatic 11.3°C ± 0.5 increase when adding 1 mM L-Asp in the presence of 10 mM Na^+^ (**Figure 2e**,**g; Supplementary Figure 6b**). Thus, L-Asp and Na^+^ bind to Anc^Na^ cooperatively, as reported for Na^+^-coupled transporters (*1, 5–8, 53, 54*). In contrast, adding L-Asp to Anc^Int^ in the absence of Na^+^ increased *T*_*m*_ by 6.9ºC ± 0.6, suggesting that, as expected, L-Asp can bind without Na^+^ (**Figure 2f-g, Supplementary Figure 6c**). Notably, 10 mM Na^+^ also stabilized the protein by 5.1ºC ± 0.6 without L-Asp and by 13.5ºC ± 0.6 in the presence of 1 mM L-Asp, suggesting that the ions can still bind to the transporter regardless of substrate binding (**Figure 2f**,**h; Supplementary Figure 6c**).

These results show that aspartate and Na^+^ bind to Anc^Int^ independently. Therefore, the intermediate Ext^Int^ clade functionally diverged from the Na^+^-coupled Ext^Na^ transporters by uncoupling substrate from Na^+^-binding. We attempted to verify that the reconstructed Anc^Int^ was robust to phylogenetic uncertainty using ‘AltALL’ sequences where all ambiguous residues are varied (Eick et al. 2017), but these produced misfolded proteins. Instead, we validated the observed functional divergence by examining the necessary and sufficient sequence changes (see below).

### *The ancestral transporter accesses high-affinity states without* Na^+^

To visualize how Na^+^-independent aspartate binding occurs, we purified Anc^Int^, reconstituted it into MSP1E3 lipid nanodiscs (**Supplementary Figure 7**), exchanged it into a buffer containing no Na^+^ or any other alkali cations (20 mM HEPES pH 7.4, 100 mM NMDG-Cl), and froze cryo-EM grids after and before adding 1 mM L-Asp, i.e., in bound and apo conditions, respectively (**Supplementary Figure 8-9**). As a Na^+^-coupled control, we reconstituted Glt_Ph_ into MSP1E3 lipid nanodiscs and froze grids under apo conditions (**Supplementary Figure 10**).

First, we compared the structures of Glt_Ph_ and Anc^Int^ under apo conditions. Cryo-EM imaging of Glt_Ph_ revealed the presence of two structural classes with final maps at 2.7 and 3.1 Å resolution (**Supplementary Figure 10-12, Supplementary Table 1**). A highly conserved non-helical loop N310-M311-D312 (NMD) motif, which breaks TM7 into TM7a and b, is the nexus for solute binding and participates in coordinating Na1, 2, and 3, as well as the substrate through its side chain and main chain atoms. In both classes, the NMD motif is in a “flipped-out” configuration, which distorts substrate- and Na^+^-binding sites (**Figure 3a,e, Supplementary Figure 13**). We call these conformations, characteristic of apo Na^+^-coupled transporters, “low-affinity” states (*11, 15, 16, 30, 55*). The two apo Glt_Ph_ structural classes differ in the position of the transport domain. One corresponds to the outward-facing state (**Figure 3a,e**) and the other to a previously observed intermediate (*16, 21, 30, 56*), where the transport domain moves toward the inward-facing state by about a quarter of the way (**Supplementary Figure 12, 13**). The HP2 gate is open in the outward-facing states, propped by M311, and closed in in the intermediate, with M311 pointing out into the lipid from underneath (**Supplementary Figure 13**).

Previously determined Glt_Ph_ and Glt_Tk_ structures show that upon Na^+^-binding to Na1 and Na3, Na^+^-coupled transporters restructure from the low-affinity apo state to a “high-affinity” state through allosterically coupled changes (*6, 7, 11, 28*) (**Figure 3b,f**). Their NMD motifs go from the ‘flipped-out’ to the ‘flipped-in’ configuration, repositioning M311 proximal to the substrate (**Figure 3b**) and altering the tilt of TM7b (*30*). R397 moves from occupying the substrate-binding site to forming a cation-π interaction with Y317 in TM7b (**Figure 3b**), its guanidium group in place to salt-bridge with L-Asp. Also, N401 repositions slightly to coordinate L-Asp optimally (*30*). Together, these changes underlie high-affinity binding by optimally positioning residues coordinating the substrate and the additional Na2 (**Figure 3c-d,g**). Following substrate binding, HP2 closes (**Figure 3g**), facilitating translocation of the transport domain to the inward-facing state.

Cryo-EM imaging of apo Anc^Int^ revealed three outward-facing structural classes, all refined to 3.0 Å resolution (**Supplementary Figure 9, 11-12, Supplementary Table 1**). One class resembles the low-affinity apo Glt_Ph_ with a ‘flipped-out’ NMD motif and an empty binding pocket; however, its R397_369_ is already in place to coordinate L-Asp (hereafter, regular text and subscript correspond to Glt_Ph_ and Anc^Int^ numbering, respectively) (**Figure 3h**). The Na^+^-binding sites are distorted similarly to apo Na^+^-coupled transporters (**Figure 3l**), except T92_78_ and [T/S]93_79_ in the Na3 site are already correctly positioned.

However, unlike apo Glt_Ph_, we also observed two apo Anc^Int^ classes in “high-affinity” states, with substrate- and Na^+^-binding sites’ architectures like those of Glt_Ph_ after Na^+^ binding to Na1 and Na3 sites (**Figure 3i,m**). The NMD motifs are flipped in, R397_369_ and N401_373_ are correctly oriented for L-Asp coordination (**Figure 3i**), and all residues in Na1 and Na3 sites are configured like in Na^+^-bound Glt_Ph_ (**Figure 3m**). Water molecules likely occupy the sites in the absence of Na^+^, but the resolution is insufficient to model them, and computational studies will be needed to understand how they compensate for the missing ions. One of the classes shows an HP2 structurally identical to substrate-bound Anc^Int^, except with an unresolved tip – the loop connecting the two helical arms of the hairpin – and an empty substrate-binding pocket (**Supplementary Figure 13**). In the other class, an ambiguous density occupies the pocket that is too large for L-Asp and might correspond to a spuriously bound buffer component; this site might be generally attractive to anions (*28*) (**Supplementary Figure 13**). We excluded this structure and poorly resolved inward-facing classes (**Supplementary Figure 12**) from further consideration.

### Binding sites are unaltered in uncoupled transporters

Imaging of aspartate-bound Anc^Int^ in Na^+^-free conditions revealed a single outward-facing structural class, which we refined to 3.4 Å overall resolution (**Supplementary Figure 8, 11-12, Supplementary Table 1**). The only structural differences between aspartate-bound Anc^Int^ and high-affinity apo Anc^Int^ are that its HP2 tip is well-structured and closed, and the substrate-binding pocket is occupied by a strong non-protein density, which we modeled as L-Asp (**Figure 3j-k,n; Supplementary Figure 13**). The substrate orientation and binding site architecture of Anc^Int^ in the absence of alkali cations were nearly identical to Na^+^/aspartate-bound Glt_Ph_ (**Figure 3d,k**). The NMD motif is “flipped-in” with the M311_289_ side chain pointing into the substrate-binding pocket and forming van der Waals interactions with L-Asp, which is coordinated by side chains of conserved T314_292_, D394_366_, R397_369_, and N401_373_ (**Figure 3k**). R397_369_ is the principal residue interacting with the side chain carboxylate of L-Asp, and D394_366_ coordinates the amino group; these residues determine the specificity of the transporter for the acidic over neutral amino acids and dicarboxylates, respectively (*57*–*59*). N401_373_, one of the most conserved residues throughout the evolution of glutamate transporters, coordinates the main chain carboxylate of L-Asp. At the current resolution, the architectures of all three Na^+^ binding sites in Anc^Int^ are identical to Na^+^/aspartate-bound Glt_Ph_ (**Figure 3n**), though Na^+^ is absent.

Our results suggest that apo Anc^Int^ spontaneously populates the high-affinity state with the correctly formed substrate- and ion-binding sites, facilitating substrate binding without Na^+^ binding. In contrast, this is a high-energy state in Glt_Ph_ and other Na^+^-coupled transporters, only accessible upon Na^+^ binding.

### Allosteric mutations turn coupling off and on

The amino acid sequence changes underlying the transition from Na^+^ coupling in Anc^Na^ to independence in Anc^Int^ are allosteric and lie outside the ion-binding sites. Anc^Na^ and Anc^Int^ are 78% identical over the transport domain sequence (**Supplementary Figure 4c**), with 26 amino acid differences having PPs above 0.9 in at least one of the ML sequences (**Supplementary Figure 4d**). The changes, however, are subtle, making it difficult to pinpoint those responsible for gaining independence from Na^+^ ions. Our phylogenetic tree shows that the evolution from Na^+^-coupled to uncoupled function occurred via several ancestral intermediates between Anc^Na^ and Anc^Int^, which might have gradually acquired the mutations. To narrow down the search for changes necessary for the functional switch, we examined a present-day transporter, Ext* (GenBank KJS87745.1), which is most homologous to the ancestor immediately preceding Anc^Int^, Anc* (**Figure 1d**). This transporter showed robust Na^+^-coupled uptake, suggesting Anc* is Na^+^-coupled (**Figure 4a**).

There are only two changes with high PPs between Anc* and Anc^Int^ transport domains - S295_273_P (0.969) and N327_305_Q (0.996), at the beginning and end of TM7, respectively (**Figure 4b**). These mutations must be sufficient to turn coupling off, and the reverse mutations in Anc^Int^ should switch coupling back on. Indeed, purified Anc^Int^ P295_273_S/Q327_305_N mutant showed Na^+^-dependent concentrative L-Asp transport in proteoliposomes (**Figure 4c**). In the thermal shift assays, L-Asp only stabilized the mutant when added with Na^+^ ions but not without (**Figure 4d; Supplementary Figure 14**). Thus, these mutations rescued Na^+^ coupling in Anc^Int^ by restoring coupled substrate and Na^+^ binding.

We compared the low-affinity and high-affinity states of Na^+^-coupled Glt_Ph_ (**Figure 4e-h**) and uncoupled Anc^Int^ (**Figure 4i-l**) to understand how allosteric mutations might affect their equilibrium. TM7 runs through the core of the transport domain with the NMD motif at its center (**Figure 4b,e,i**). In Na^+^-coupled transporters Glt_Ph_ and human EAAT3, TM7b shows a significantly different tilt in the low-affinity apo state compared to the high-affinity Na^+^-bound state (**Figure 4e,f**), translating into repacking of surrounding TMs3, 6, 8, and HP2 (**Figure 4g-h**). Packing differences in this region affect Glt_Ph_ affinity for L-Asp and are, therefore, allosterically coupled to the changes at the binding site (*3, 60, 61*). In contrast, TM7b of Anc^Int^ shows little tilt changes and less helical repacking in the low-affinity state compared to the high-affinity state (**Figure 4i-k**), suggesting that S295_273_P and N327_305_Q substitutions stabilize the high-affinity state. P295_273_ is in the loop preceding TM7a and might rigidify the loop compared to S (**Figure 4b,i**). Q327_305_ is the last residue of a highly conserved F-[I/V]-A-[N/Q] signature sequence motif in TM7b (*62*) (**Figure 4b,i**). Q327_305_ hydrogen bonds to the top of TM3 (**Figure 4l**), perhaps stabilizing TM7-TM3 interactions. By contrast, N327_305_ in Glt_Ph_ is too short to reach TM3 (**Figure 4h**). These analyses suggest that the structural flexibility of the central TM7 determines the equilibrium between the transporter states, and a more rigid TM7 in Anc^Int^ tightly packed to surrounding helices might eliminate the need for Na^+^-binding to achieve the high-affinity state.

## Discussion

Na^+^-coupled glutamate transport is one of the most thoroughly characterized ion-coupled transport mechanisms, where Na^+^ binding is inextricably linked to substrate binding. Yet, how these binding sites are allosterically coupled and, thus, the design principles of Na^+^-coupled transporters have remained unknown. We gained unprecedented insights into this fundamental question by examining the evolution of prokaryotic glutamate transporters using phylogenetics and APR. Surprisingly, the transition from powering transport with Na^+^ to H^+^ gradients occurred via an intermediate ancestral transporter Anc^Int^, in which aspartate binding was ‘uncoupled’ from Na^+^ binding. It remains unclear why Anc^Int^ showed no transport in proteoliposomes. We observed outward- and inward-facing structural classes in cryo-EM images, suggesting that Anc^Int^ could undergo conformational transitions necessary for transport. Ancestral proteins are generally more thermostable than extant transporters for evolutionary and methodological reasons (*4, 63*), which could manifest as slow transitional kinetics, perhaps making transport too slow to observe.

Unlike Na^+^-coupled homologs, Anc^Int^ did not require Na^+^ to form high-affinity substrate- and ion-binding sites. Thus, we propose that the relative free energies of the low- and high-affinity states are the linchpin of coupling (**Figure 5a-c**). In Na^+^-coupled transporters, the free energy of the low-affinity conformation is much lower than the high-affinity conformation (**Figure 5b**). The substrate binding energy alone is insufficient to overcome the penalty, and Na^+^ binding pays the energetic price, stabilizing the state. In uncoupled Anc^Int^, the low-affinity state has a similar or higher energy, and the high-affinity state is populated without ions (**Figure 5c**). While Na^+^ ions can still bind, they are unnecessary for substrate binding.

We found two high-probability allosteric mutations between uncoupled Anc^Int^ and the previous ancestor inferred to be Na^+^-coupled. In Anc^Int^, these were sufficient to restore Na^+^ coupled binding and transport, likely by stabilizing the low-affinity state. However, it seems likely that before this switch, transporters accumulated other neutral epistatic mutations during the evolution from Na^+^-coupled Anc^Na^ to Anc^Int^, which gradually increased the relative energy of the low-affinity state. Then, the last two changes were sufficient to make the high-affinity state accessible and function independent of Na^+^ (**Figure 5c**). Such allosteric alterations of energy landscapes might underlie the evolutionary diversification of many proteins (*64*–*67*). The mechanism explains how a few mutations can turn off Na^+^ coupling without eliminating the three Na^+^-binding sites through multiple mutations, which would diminish the transporter’s ability to bind substrate and fitness.

Was the transition to uncoupled substrate binding a random evolutionary event or an adaptation required for organismal fitness? While functional studies of Anc^Int^ descendants will need to be carried out, most are unlikely to be ion-coupled based on a lack of Na^+^ or H^+^-coupled signature residues. Transporter copies or transporters from different families with similar functions could compensate for the loss of coupled transport (*68*), which still begs the question of uncoupled transporters’ physiological relevance. Examining the gene neighbors of uncoupled transporters revealed that many are adjacent to enzymes involved in amino acid metabolism; we observed no similar preferences in the Na^+^- or H^+^-coupled gene neighbors (**Supplementary Figure 15**). Thus, uncoupled transporters might mediate amino acid uptake paired with their enzymatic utilization. Such pairing would be reminiscent of glucose transporters, which pair uncoupled transport with subsequent glucose metabolism, a key interplay for insulin secretion (*69*).

In summary, we show that phylogenetic analysis and APR can uncover the evolutionary trajectory of functional diversification in transporters and pinpoint amino acid changes that alter the function. Comparative studies of ion-coupled and -independent proteins show that coupling can be allosterically regulated and turned on and off by mutations and perhaps environmental changes, which reshape the energy landscape of the transporter. We suggest that drugs could also allosterically modulate the extent of ion and substrate coupling in secondary active transporters, allowing novel inhibitory modality in which transport is not abolished but its concentrative capacity is diminished. Such drugs could offer advantages in cases where complete inhibition might be harmful.

## Methods

### Sequence retrieval, curation, and alignment

We designed a workflow to construct maximum likelihood phylogenies capturing the Na^+^-coupled to H^+^-coupled (Ext^Na^ to Ext^H^) evolutionary transition based on previous APR studies (*47*) (**Supplementary Figure 2**). Biochemically characterized Ext^H^ proteins feature signature sequence substitutions in the Na^+^-binding sites compared to Ext^Na^ transporters (*2, 25, 70*); [T/S]92[A/L/I/V/M], D405N, and M311L (numbering is based on Ext^Na^ archaeal transporter Glt_Ph_) (**Figure 1c**). Putative Ext^H^ genes carrying these signature residues are widespread throughout bacterial phyla. We constructed a maximum parsimony tree of all bacterial glutamate transporter homologs and found that putative Ext^H^ transporters cluster together regardless of phylum (**Supplementary Figure 3a**). Though we cannot exclude the existence of H^+^-coupled transporters with completely different signature residues, we infer that the emergence of Ext^H^ was a single gene duplication event, following which Ext^H^ genes spread through horizontal gene transfer.

Aiming to have a sequence set with a size amenable to APR, we retrieved 20,000 non-redundant *Bacillota/Firmicutes* sequences via NCBI PSI-BLAST since Ext^H^ sequences were predominantly found in this phylum. We used WP_011876993.1 as the original query (*71, 72*). After clustering by 99% using CD-HIT (*73*), removal of sequences with ambiguous amino acid assignment, and alignment by MUSCLE v5 (*74*), a maximum parsimony tree was generated by MP-boot (*75*). Subsets of sequences were selected based on proximity to sequences with signatures of H^+^-coupling and nearby sequences up until those with signature Ext^Na^ residues. The remaining sequences were clustered to 90% using CD-HIT, and the resulting alignment was manually curated to exclude sequences with major gaps or insertions. Archaeal Glt_Ph_ (UniProt ID O59010) was included to be the tree root. The final sequence set (1,153 sequences) was aligned by MAFFT-DASH, which incorporates information from PDB structures into the alignment matrix (*76*). Finally, variable loops and tails were removed, and the resulting alignment was manually adjusted to account for poorly aligned regions.

### Phylogenetic analysis, ancestral sequence reconstruction, and construct design

Maximum-likelihood (ML) trees and ancestral sequences were generated using IQ-TREE 2.1.4-beta (*77*). Evaluation of all potential evolution models and rate heterogeneity parameters using ModelFinder revealed that LG+F+R was the best protein evolution model (*78*–*80*). FreeRate (F) heterogeneity considers site-specific differences in evolutionary rate, relaxing the constraint of equal site probabilities imposed by discrete Gamma rate heterogeneity. The number of possible rate categories that could be assigned to a site (R) was empirically determined for each run using ModelFinder to query a range of 10-20.

As with other APR studies of microbial proteins, our tree topology is not reconciled to the microbial species tree, as this has poor confidence especially at lower taxonomic ranks (*81*). We chose the outgroup-based approach of rooting the tree, using archaeal Glt_Ph_ as the root due to the evolutionary distance between bacteria and archaea. Due to the difficulties of reconciling microbial gene trees to species trees, our phylogeny might not reflect the true chronology of evolutionary history (*81*). While Na^+^-coupled transporters appear most ancient (**Supplementary Figure 3b**), alternate rooting, placing Ext^Int^ or Ext^H^ as the most ancient, is possible (**Supplementary Figure 3c**) but unlikely because most archaeal transporters contain Na^+^ coupling signature residues.

After performing over 60 replicate tree runs, we found that the best *Firmicutes* tree yielded robust UFBoot2 values, ranging between 91-100% for the Ext^H^ node (**Figure 1d**). When we repeated the analysis for the *Proteobacteria* phylum, the best tree had 5-87% UFBoot2 values for the equivalent nodes, reflecting non-reproducible trees. Thus, we used the *Firmicutes* tree for all subsequent analyses.

The run yielding the best tree, as determined by log-likelihood, was inspected for long branches, defined as > 0.7 substitutions/site. These were removed; in the second round of tree-searching, the analysis was repeated with 254 independent tree search runs. The final tree had the best-fit model of LG+F+R14, chosen according to the Bayesian Information Criterion. Bootstrap values were determined by UFBoot2 using 1000 replicates (*45*). Ancestral sequences were reconstructed from the best tree using the IQ-TREE implementation (*77*) based on Yang’s marginal reconstruction method (*82*–*84*). The ML sequences were generated using custom Python scripts, either written manually or generated using ChatGPT. The final ML amino acid sequence for each node is constructed using the amino acid with the highest posterior probability (PP) at every site. All tree figures were generated using TreeViewer (*85*). Visual representations of sequence alignments and posterior probabilities were made using WebLogos (*86*).

### Protein expression and purification

Since poorly aligned loops and tails are not reconstructed in our APR analysis, we had to graft them in for expression constructs. We took comparable loops and tails from an extant sequence close to Anc^Int^ (KJS87745.1); the 3L4 has two lysines that were mutated to histidines (K105H, K109H) to prevent potential proteolysis. All protein constructs were codon-optimized for *E. coli*, synthesized (GenScript), and cloned into a modified pBAD vector containing a C-terminal thrombin cleavage site followed by either an ALFA peptide tag (*87*) or a C-terminal thermostable GFP which is specialized for membrane proteins (*49*). An additional Twin-Strep tag and a 10x-HIS tag were added to the C-terminus of these constructs, all separated by glycine-serine (GSSS) linkers. The Glt_Ph_ construct used was the ‘CAT7’ construct, with seven histidines previously introduced for stability (*13*). Vectors were transformed into *E. coli* BL21(DE3) or DH10B cells, cells were grown in 2xYT media, and expression was induced by adding 0.1% arabinose at 0.8-1.0 OD_600_. Proteins were expressed for 2-3 hours at 37°C. Membranes were purified as previously described for Glt_Ph_, with all steps being performed at 4°C (*3*). Briefly, cells were broken using a high-pressure cell disruptor (Avestin) for three passes with cooled water continuously applied. The resulting lysate was spun at 17,400 x g for 15 min, and the supernatant was spun at 186,000 x g for 1 hour. The resulting membrane pellet was homogenized in 20 mM HEPES/NaOH pH 7.4, 10 mM EDTA, and 10% Sucrose; finally, the slow and fast spins were repeated, and the resulting pellet was frozen at -80°C.

All protein purification steps were performed at 4-8°C. Crude membranes were homogenized in 20 mM HEPES/NaOH, pH 7.4, 200 mM NaCl, 1 mM monopotassium L-aspartate (L-Asp); solubilization was initiated by the addition of 40 mM n-dodecyl-β-D-maltopyranoside (DDM; Anatrace), and the solubilizing membranes were gently rocked for two hours. Over the course of our experiments, we noticed that tags were being non-specifically proteolyzed upon solubilization; to reduce this, we added 1 mM PMSF and 1 mM EDTA before adding DDM. After solubilization, samples were clarified by high-speed ultracentrifugation at 186,000 x g for 1 hour.

For ancestral constructs, the supernatant was applied to a prepacked 1 mL Streptactin XT 4Flow high-capacity column (IBA Biosciences) at a constant flow rate of 0.5-1 mL/min. The resin was washed until UV_280_ absorbance was flat, corresponding to fourteen column volumes of 20 mM HEPES/NaOH, pH 7.4, 200 mM NaCl, 1 mM L-Asp, and 1 mM DDM. Subsequently, the resin was eluted with the same buffer containing 50 mM biotin (IBA Biosciences), and proteins were concentrated.

For Glt_Ph_, the supernatant was applied to a 1 mL HisTrap HP (Cytiva) at a constant flow rate of 1 mL/min. The resin was washed until UV_280_ absorbance was flat, corresponding to thirty column volumes of 20 mM HEPES/NaOH, pH 7.4, 200 mM NaCl, 1 mM L-Asp, 40 mM imidazole, and 1 mM DDM. Subsequently, the resin was eluted with the same buffer containing increased imidazole (250 mM), and proteins were concentrated.

### Reconstitution into proteoliposomes

Unilamellar liposomes were prepared essentially as previously described (*3*). Briefly, a 3:1 (w:w) ratio of *E. coli* polar lipids and egg phosphatidylcholine (Avanti Polar Lipids) was dried, hydrated in 50 mM HEPES/Tris, pH 7.4, 200 mM KCl at a final concentration of 5 mg/mL, flash frozen, and stored at -80°C. On the day of the reconstitution, liposomes were thawed, extruded through a 400 nm filter 13 times, and destabilized with Triton X-100 using a 1:2 (w:w) Triton X-100 to lipid ratio. For the counterflow and Na^+^-coupled uptake experiments, GFP-free proteins were purified by SEC using Superdex 200 Increase 10/300 column (Cytiva) in 20 mM HEPES/NaOH, pH 7.4, 200 mM NaCl, 1 mM L-Asp, and 1 mM DDM; proteins were subsequently concentrated using a 100 kDa cutoff concentrator. To form proteoliposomes, the purified protein was mixed with destabilized liposomes (1:100, w:w) for 30 minutes, then detergent was removed with BioBeads SM-2 (Bio-Rad) at 80 mg/ml, which were applied twice at 22°C for two hours, once overnight at 4°C, and once more at 22°C for two hours.

### Proteoliposome uptake experiments

After detergent removal with BioBeads, proteoliposomes underwent the same general procedure for buffer exchange and concentration, regardless of experiment. Proteoliposomes were exchanged into the appropriate buffer using a PD-10 column (Cytiva), flash frozen in liquid N_2_, thawed, concentrated via ultracentrifugation (100,000 x g for one hour), resuspended to a final concentration of 50 mg/mL, flash frozen, and stored at -80°C. On the day of the experiment, the proteoliposomes were thawed and extruded through a 400 nm filter 23 times. To minimize sample dilution in the extruder, the proteoliposome volume was no less than 100 μL. All subsequent experiments were performed at 30°C.

For uptake experiments, liposomes were prepared as described above to a final concentration of 50 mg/mL in 20 mM HEPES/Tris pH 7.4, 200 mM KCl, and 100 mM choline chloride (ChoCl). To initiate Na^+^-driven uptake in the presence of a negative membrane potential (-102 mV), liposomes were diluted 1:100 in 20 mM HEPES/Tris pH 7.4, 2 mM KCl, 198 mM choline chloride, 100 mM NaCl, 0.9 μM valinomycin, and 1 μM L-aspartate. The L-aspartate was a mixture of cold monopotassium L-aspartate (RPI) and ^3^H-L-Asp (PerkinElmer or American Radiolabeled Chemicals). For negative controls without Na^+^-gradient, the NaCl was replaced with choline chloride.

To stop the reactions, 200 μL of reaction mixture (100 μg of liposomes) was diluted in 2 mL ice-cold quenching buffer (20 mM HEPES/Tris pH 7.4, 200 mM lithium chloride). The quenched reaction was immediately filtered through a 0.22-μm filter membrane (Millipore Sigma) and washed three times with 2 ml quenching buffer. Washed membranes were inserted into scintillation vials, and the membranes were soaked overnight in 5 ml Econo-Safe Counting Cocktail. Radioactivity in liposomes was measured using an LS6500 scintillation counter (Beckman Coulter). Technical replicates from an individual reconstitution were taken in duplicate and averaged, and at least two separate reconstitutions were taken per point.

### Nano Differential Scanning Fluorimetry (nanoDSF)

All steps were performed on the same day of the protein solubilization and purification, since we observed variance in results depending on the freshness of the protein. We opted to use GFP-tagged ancestral proteins because the *T*_*m*_ of the fused GFP is greater than 90ºC (*49*), which does not interfere the *T*_*m*_s observed in our experiments. After affinity purification, the proteins with C-terminal GFP were concentrated using a 100 kDa cutoff concentrator (MilliporeSigma). To remove ligands, the protein was dialyzed using a PD MiniTrap G-25 (Cytiva). For Anc^Na^ and Anc^Int^ P295_273_S/Q327_305_N, this buffer was 20 mM HEPES, 200 mM NMDG-Cl, pH 7.4, and 1 mM DDM; for Anc^Int^, this buffer was 50 mM KH_2_PO_4_/K_2_HPO_4_ buffer, pH 7, and 1 mM DDM. Different buffers were used for optimal first derivative curves in apo conditions, and we observed identical *T*_*m*_s in substrate-bound conditions regardless of dialysis buffer composition. The dialyzed, apo protein was concentrated and diluted into the same respective buffers at a final concentration of ∼1 mg/mL; for Anc^Int^, the buffer contained 10 mM decyl-maltoside (DM) instead of 1 mM DDM. This was necessary to observe melting of Anc^Int^, presumably by the shorter chain detergent destabilizing the overall protein. Thermal stability in the presence and absence of various potential ligands was measured using a Tycho NT.6 instrument (Nanotemper), and individual *T*_*m*_s were calculated by determining the appropriate local minimum in the first-derivative of the 350/330 nm ratio.

### Microscale thermophoresis (MST)

After affinity purification, the proteins with C-terminal GFP were concentrated using a 100 kDa cutoff concentrator, then SEC-purified on Superose 6 Increase 10/300 column (Cytiva) in 50 mM KH_2_PO_4_/K_2_HPO_4_ buffer pH 7 and 1 mM DDM. For the MST experiment, monopotassium L-aspartate was serially diluted 2-fold in water for a total of 16 concentrations. Each aspartate dilution was mixed with in a 1:1 (v:v) ratio of protein samples, so that the final protein concentration was 100-300 nM, the appropriate salt was 100 mM, the DDM was 0.4 mM, and the KH_2_PO_4_/K_2_HPO_4_ was 10-20 mM. Measurements were taken using Monolith NT.115 Blue/Red MST (Nanotemper), High MST power, Blue detector (for GFP), and premium capillaries (Nanotemper).

For data analysis, obvious outliers caused by aggregates (as determined by instrument software) were discarded. The ideal response time for each individual experiment was determined by instrument software, and this was always between 1.5s-5s. Individual binding titrations were normalized to fraction bound by *K*_*D*_ fits in the instrument software. To obtain the final binding affinity value, the experiments were repeated three independent times, the data points were averaged, and the *K*_*D*_ was calculated from the average values.

### Cryo-EM sample preparation

Membrane Scaffold Protein (MSP1E3) was expressed and purified as previously described (*88*) in the absence of Na^+^ or L-Asp. A 3:1 (w:w) ratio of *E. coli* polar lipids and egg phosphatidylcholine (Avanti Polar Lipids) was dried, hydrated in 50 mM HEPES/Tris, pH 7.4 and 200 mM KCl, and solubilized in 72 mM DDM at final lipid concentration of 18 mM as previously described (*15, 88*).

For Anc^Int^, GFP-free transporter constructs were affinity purified, concentrated, and mixed with MSP1E3 and solubilized lipids (both containing no Na^+^ or L-aspartate) at a molar ratio of 1.5:1:50, aiming for a final concentration of lipid greater than 5 mM. This mixture was incubated at 22°C for 30 minutes, then 350 mg BioBeads SM-2 (BioRad) per mL mixture (w:v) was added for two hours at 22°C with gentle shaking, and another round of BioBeads was applied overnight at 4°C with gentle shaking. The resulting MSP1E3/protein preparations were cleared by ultracentrifugation at 100,000 x g for 30 minutes. The supernatant was applied to a Superose 6 Increase 10/300 column in 20 mM HEPES pH 7.4, 100 mM NMDG-Cl, and the peak fractions were concentrated to 7-9 mg/mL, as measured by A280 absorbance. For the Anc^Int^ substrate-bound dataset, 1 mM L-Asp was added to the sample just before freezing. Reconstitution efficiency was verified using SDS-PAGE.

Glt_Ph_ was reconstituted into MSP1E3 as previously described (*15*). Affinity purified protein was further purified via size-exclusion chromatography (Superdex 200 Increase, 10/300). Peak fractions were concentrated, mixed with MSP1E3 and solubilized lipids at a molar ratio of 0.75:1:50, aiming for a final concentration of lipid greater than 5 mM. This mixture was incubated and treated with BioBeads SM-2 in the exact same fashion as ancestral protein. The resulting MSP1E3/protein preparations were exchanged in Na^+^-free buffer by three rounds of 10-fold dilution/concentration, and cleared by ultracentrifugation at 100,000 x g for 30 minutes. The supernatant was applied to a Superose 6 Increase 10/300 column in 20 mM HEPES/Tris pH 7.4 and 50-mM choline chloride, with the goal of mimicking previously published conditions. The peak fractions were concentrated to 8 mg/mL, as measured by A280 absorbance. Reconstitution efficiency was verified using SDS-PAGE.

To improve particle distribution, fluorinated Fos-Choline-8 (Anatrace) dissolved in water was added to the sample to a final concentration of 1.5 mM. UltrAuFoil R1.2/1.3 300-mesh gold grids (Quantifoil) were glow-discharged at 25 mA for 80 s. 3 μL of protein was applied to each grid, incubated for 20 s under 100% humidity at 4°C, blotted for 2.5 s under 0 blot force, and plunge frozen in liquid ethane using a Vitrobot Mark IV (Thermo Fisher Scientific).

### Cryo-EM data acquisition and processing

Imaging was performed on a Titan Krios (FEI) equipped with a K3 camera (Gatan) using Leginon software. Specifics of each data collection and each individual map and model are provided in **Supplementary Table 1**. Visual summaries of processing workflows and EM validation are provided in **Supplementary Figures 8-11**.

### Initial processing pipeline for Anc^Int^

For all datasets, raw movies were imported into cryoSPARC 4.0-4.4 and subjected to Patch Motion Correction and Patch CTF Estimation, where the movies were binned 2x (*89*). Low-quality movies were removed, as determined by CTF fits, ice thickness, intensity, etc. Particles were manually picked to train Topaz (*90*), where for each data set, the optimal picking model and estimated number of particles per micrograph were empirically determined using the Topaz Cross Validation tool in cryoSPARC. After picking and extraction with 4x binning, an *ab initio* 3D model was constructed, and high-quality particles were sorted using multiple heterogeneous refinement jobs, where ‘good’ particles sort into the *ab initio* model class, and ‘junk’ particles sort into three pure noise ‘decoy’ volume classes (created by performing one iteration of *ab initio*). When greater than 95% of particles converged into the protein volume, the remaining particles were re-extracted to full box size and subjected to 3D Classification without alignment, and Non-Uniform (NU) Refinement (*91*). To maximize the number of particles obtained, the Topaz model was retrained on the clean particle stack after rounds of 2D classification, and heterogeneous refinement was repeated with the new particle stack and the high-resolution volume from NU-Refinement (replacing the *ab initio* volume). Remaining particles were re-extracted to full box size and subjected to NU-Refinement.

### Anc^Int^ apo data set processing

Our ancestral protein datasets were highly heterogeneous and poorly symmetric; throughout the processing we realized that they were reminiscent of other seemingly mobile membrane protein structures (*92, 93*). To overcome this, we attempted to enrich particles with symmetric features so that we could classify individual protomers. In the apo dataset, we converted particle coordinates to Relion format using PyEM (*94*). Particles were re-extracted in Relion using movies motion corrected and binned 2x using Relion’s implementation of MotionCor2 (*95*). These particles were re-imported into cryoSPARC, refined using NU-Refinement (C1), and the resulting particle images were converted to Relion format with PyEM. These particles were then subjected to two rounds of Bayesian polishing (*96*), classification with heterogeneous refinement, and classification with NU-Refinement (C1). To remove ‘disordered’ particles, we used 3D classification without alignment (C1) in cryoSPARC. We further optimized symmetric particles by performing one round of *ab initio* with C3 symmetry imposed. Using the best class, we achieved a resolution of 2.8 Å in C3, which allowed us to resolve individual protomers using symmetry expansion and focused classification using a mask on the transport domain. Each class underwent local refinement masked on the entire protomer (including transport/scaffold domains) using pose/shift Gaussian priors during alignment (standard deviations of 3º, 2 Å).

### Anc^Int^ substrate-bound processing

As with the apo dataset, the substrate-bound dataset was similarly heterogeneous and asymmetric. Using the previous approach, we encountered trouble imposing C3 symmetry, preventing focused classification of individual protomers. During our processing, we realized that we could more efficiently remove ‘disordered’ particles using outputs from 3D Variability, where the first variability component reflects this ‘order-disorder’ transition between rigid, easily aligned particles and presumably mobile, difficult to align particles. We then took the first frame of the 3DVA movie (disordered, ‘bad’ volume), the last frame of the movie (ordered, ‘good’ volume), and two decoy noise volumes as inputs for heterogeneous refinement. Though a fundamentally similar approach to 3D classification used in the apo dataset, the new approach allowed us to impose C3 symmetry with good resolutions (3.4 Å). We then symmetry expanded the particles, performed 3D classification without alignment on the masked transport domain, and locally refined the protomers using pose/shift Gaussian priors during alignment (standard deviations of 3º, 2 Å), as with the apo dataset.

### Glt_Ph_ apo processing

Raw movies were imported into cryoSPARC 4.4 and subjected to Patch Motion Correction and Patch CTF Estimation, where the movies were binned 2x (*89*). Low-quality movies were removed, as determined by CTF fits, ice thickness, intensity, etc. Particles were manually picked to train Topaz (*90*), where the optimal picking model and estimated number of particles per micrograph were empirically determined using the Topaz Cross Validation tool in cryoSPARC. After picking and extraction, an *ab initio* 3D model was constructed. After one round of heterogeneous refinement with *ab initio* volumes, we performed NU-Refinement (C3) on the remaining particles, symmetry expanded the particles, performed 3D classification without alignment on the masked transport domain, and locally refined the protomers using pose/shift Gaussian priors during alignment (standard deviations of 3º, 2 Å).

### Model building and refinement

For model building, maps were sharpened using DeepEMhancer (*97*). We generated a homology model of Anc^Int^ in SWISS-MODEL, using an outward-facing state of Glt_Ph_ (PDB ID 2NWX) as a template (*13, 98*). The templates for Glt_Ph_ were either 6UWF (OFS) or 6UWL (iOFS). The model was adjusted into the density using ISOLDE (*99*), and the final model was iteratively real-spaced refined and validated in Phenix using the unsharpened density and default parameters (*100*). All structural figures were made with ChimeraX (*101*).

### Gene neighborhood analysis

For each clade (Ext^Na^, Ext^Int^, and Ext^H^), a sequence similarity network (SSN) and corresponding genome neighborhood was created using the Enzyme Function Initiative (EFI) webserver (*102*). Transporters without information regarding protein-encoding gene neighbors were excluded. A transporter was considered paired to an enzyme if they were within 3 neighboring genes away from the transporter. Genes were considered to have amino acid/dicarboxylate modifying activity based on Pfam annotation.

## Data availability

Atomic coordinates and EM densities have been deposited in PDB and EMDB for the following structures: Apo Anc^Int^ low-affinity (PDB 9BGY, EMD-44526); Apo Anc^Int^ high-affinity (PDB 9BGZ, EMD-44527); L-Asp bound Anc^Int^ (PDB 9BH0, EMD-44528); Apo Glt_Ph_ iOFS (PDB 9BH1, EMD-44529); Apo Glt_Ph_ OFS (PDB 9BH2, EMD-44530). Bioinformatic analyses, structures, and EM maps will be made available immediately upon publication.

## Author contributions

K.D.R. and O.B. conceived the project and wrote the manuscript. K.D.R. performed all evolutionary analyses. K.D.R. and B.R. performed the functional experiments and prepared the figures. K.D.R., B.R., and F.B.A. prepared grids and processed cryo-EM data. K.D.R., B.R., F.B.A., and O.B. built and refined the molecular models. N.K. and S.P. performed foundational pilot experiments, including expression/purification optimization, and preliminary cryo-EM data collection and processing. All authors reviewed and edited the manuscript.

## Acknowledgments

We thank the following the people for their contributions: Joseph Thornton and members of the Thornton lab, Michael Harms, Luce Skrabanek, and Eva Fortea for helpful discussions regarding bioinformatics and phylogenetics; members of the Boudker lab, especially Amanda Scopelliti, Xiaoyu Wang, Biao Qiu, and Yessenia Lopez, for helpful discussions regarding biochemistry and cryo-EM; Alessio Accardi and Crina Nimigean for critical reading of the manuscript; Will Eng for assistance with protein expression; GenScript for prompt and patient support with gene synthesis; Vanessa Braband, Bridget Milorey, and Dinorah Leyva from NanoTemper for helping with troubleshooting MST parameters. The work was supported by National Institutes of Health (NIH) grants F32 NS102325 (to K.D.R.) and R37NS134865 (to O.B.). O.B. is an investigator of the Howard Hughes Medical Institute.

Phylogenetic analyses were performed on the Weill Cornell HPC cluster maintained by the Scientific Computing Unit. ChatGPT was used to write some of the Python scripts involved in bioinformatic data analysis. All cryo-EM grids were initially screened at the Weill Cornell Cryo-EM Core Facility with help from Carl Fluck. Apo Anc^Int^ was collected at the NYU cryo-EM facility with help from Bing Wang. Substrate-bound Anc^Int^ was collected at the Columbia University Medical Center Cryo-EM core with assistance from Robert Grassucci. An earlier substrate-bound Anc^Int^ dataset (with assistance from Kasahun Nehelu) and the apo-Glt_Ph_ dataset (with assistance from Aaron Owji and Aygul Ishemgulova) were collected at the Simons Electron Microscopy Center and National Resource for Automated Molecular Microscopy located at the New York Structural Biology Center, supported by grants from the Simons Foundation (349247), NYSTAR, and the NIH National Institute of General Medical Sciences (GM103310) with additional support from Agouron Institute (F00316) and NIH S10 OD019994-01.

## Notes

### Competing Interest Statement

The authors have declared no competing interest.

### Summary of Updates

Additional figure (new Figure 4), updates on text and figures.

