## Supplementary File for "Evolutionary analysis reveals the origin of sodium coupling in glutamate transporters"

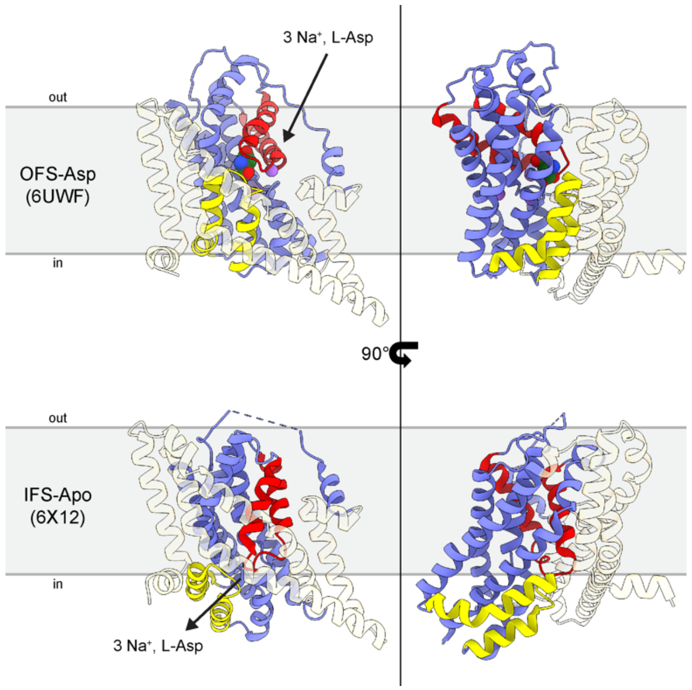


**Supplementary Figure 1. Elevator mechanism of SLC1-type transporters.** $Glt_{Ph}$ structures in the substrate-bound outward-facing state (OFS-Asp, PDB 6UWF) and substrate-free inward-facing open state (IFS-Apo, 6X12). The rigid scaffold domain is in transparent wheat; the mobile transport domain is in blue; HP1 is in yellow; HP2 is in red; purple spheres are $Na^+$; green molecule is L-Asp. In a typical transport cycle, three $Na^+$ ions and L-Asp bind in the outward-facing state, resulting in the closure of the HP2 gate and translocation of the transport domain to the inward-facing state. The HP2 gate then opens, releasing solutes to the intracellular side.

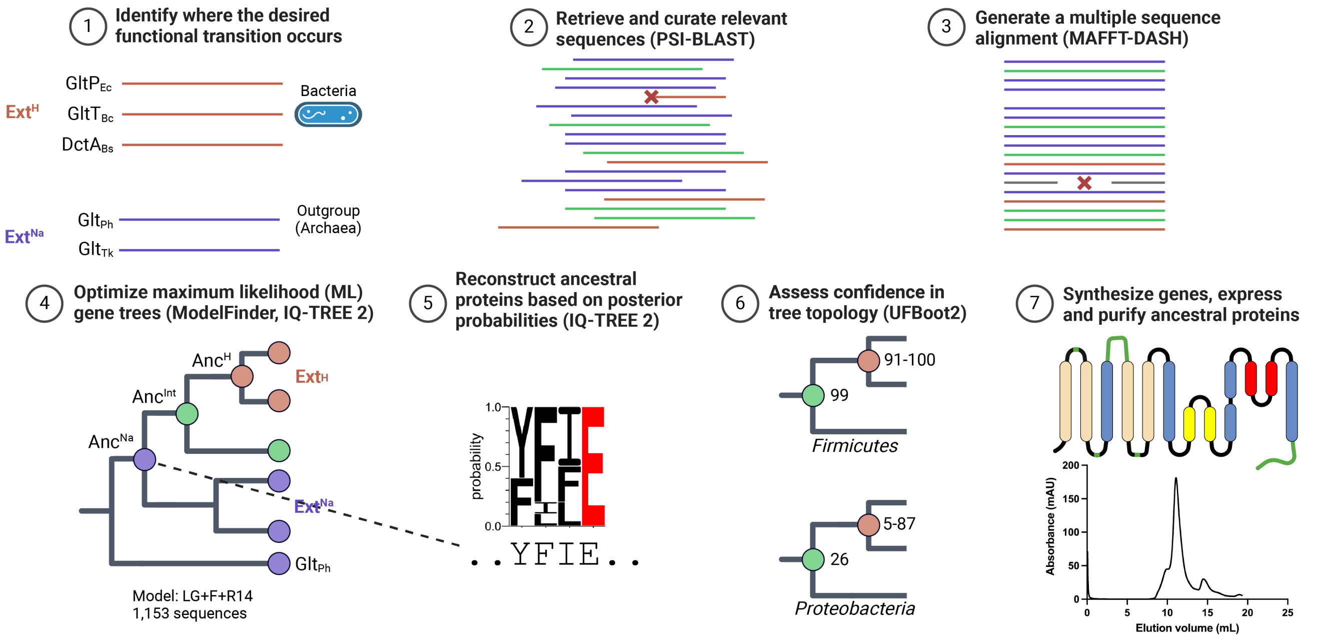


**Supplementary Figure 2. ASR workflow to capture Ext$^{Na}$ to Ext$^{H}$ evolutionary transition.** The diagram is based on Orlandi et al. and was created using BioRender. (1) Based on previous biochemical and structural characterization of prokaryotic $H^+$-coupled (orange; GltP$_{Ec}$, GltT$_{Bc}$, DctA$_{Bs}$) and $Na^+$-coupled (purple; Glt$_{Ph}$, Glt$_{Tk}$) transporters, we identified signature residues that helped us sort putative sequences as either Ext$^{H}$ or Ext$^{Na}$. (2) We retrieved homologs of transporters using PSI-BLAST. After clustering and quick sequence alignment, we removed sequences with large gaps or insertions. Finally, we used parsimony tree generation to keep sequences that encompass Ext$^{H}$, Ext$^{Na}$, and intermediate (green) transporters. (3) We carefully aligned the sequences using MAFFT-DASH, which incorporates structural information. Sequences with large gaps or insertions were removed. The remaining sequences were re-aligned, and poorly aligned loops and tails were stripped from the alignment. (4) The final sequence alignment was used as input for ML tree generation, using archaeal Glt$_{Ph}$ as the root. The optimal model of protein evolution was selected using ModelFinder. Sequences that caused long branches were removed, and 254 parallel tree searches were performed to find the optimal branch lengths and topology, resulting in the ML tree with the highest possible likelihood. (5) To reconstruct ancestral sequences, every node was reconstructed using optimized ML parameters and protein evolution model; each site in each node is assigned probabilities for all 20 amino acids, and the highest-probability amino acid at a site is selected for the ML sequence. (6) To determine if a given node is creating a robustly reconstructed ancestor, bootstrap is performed (UFBoot2), which calculates branch supports by resampling the MSA 1000 times and determining the frequency of the branch re-occurring in the trees. Sequence sampling is critical in this step; when we retrieved sequences from *Firmicutes/Bacillota*, we obtained robust UFBoot2 values for Anc$^{Int}$ and descendants, but not when retrieving sequences from *Proteobacteria*. (7) Ancestral sequences are engineered for expression and purification by inserting loops and tails from extant sequences (green), which were not considered during ASR due to poor alignment. All relevant references here and elsewhere in the legends are in the Main Text or Methods.

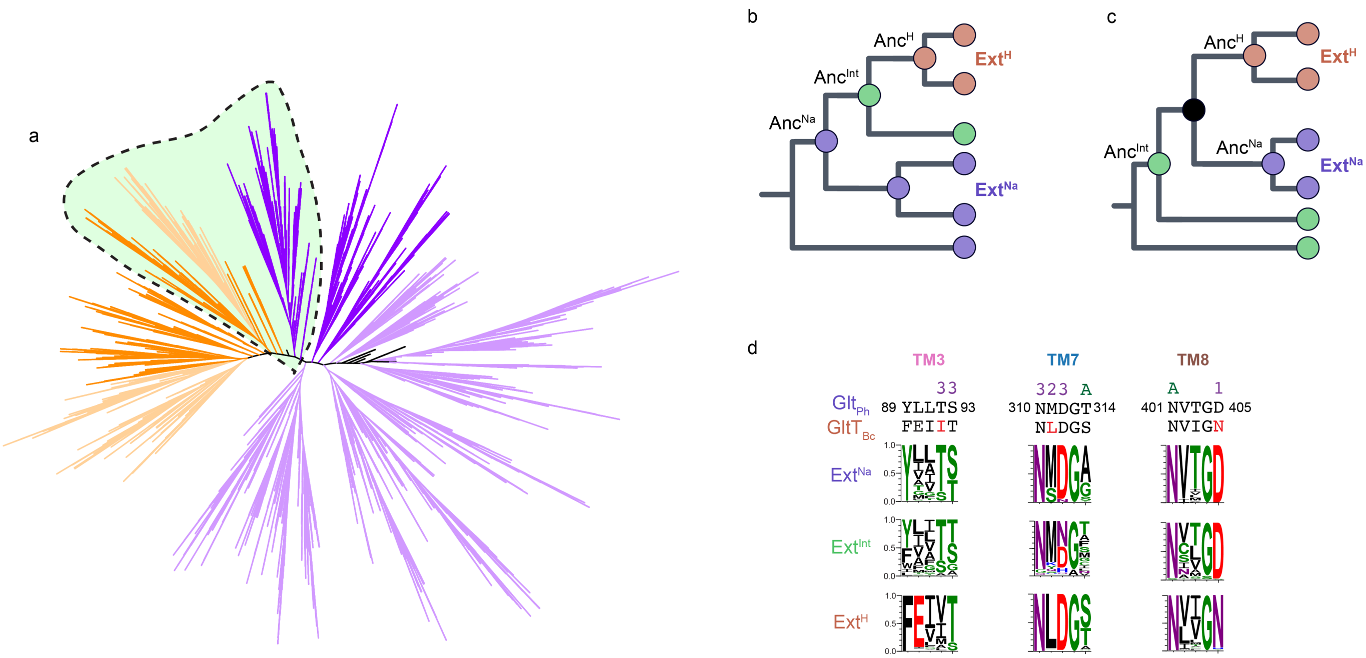


**Supplementary Figure 3. Phylogeny, rooting, and conservation of bacterial glutamate transporter homologs.** (a) Maximum parsimony phylogeny of bacterial glutamate transporter homologs. Clades containing sequences with conserved signature residues of $Ext^{Na}$ and $Ext^{H}$ transporters are dark purple and dark orange, respectively; light purple and light orange clades indicate less conservation of critical residues. The green area highlights the region used for subsequent phylogenetic analyses. (b-c) Potential rooting of $Ext^{Na}$ to $Ext^{H}$ transition. Illustrations represent the rooting to $Glt_{Ph}$ performed in this study (b) and an alternative root placed at $Anc^{Int}$ as the earliest ancestor (c). Illustrations were created in BioRender. (d) Conservation of the $Na^+$-binding sites throughout the phylogeny. Top: sequences of $Na^+$-coupled $Glt_{Ph}$ and $H^+$-coupled $Glt_{Bc}$ encompassing $Na^+$-binding sites of $Ext^{Na}$. The numbers above the sequences mark side chains in $Glt_{Ph}$ that coordinate $Na^+$ sites 1-3; A indicates side chains coordinating L-Asp. Bottom: WebLogos of the relative amino acid frequencies in $Ext^{Na}$, $Ext^{Int}$, and $Ext^{H}$ clades. The number of sequences per clade is 186, 161, and 773, respectively.

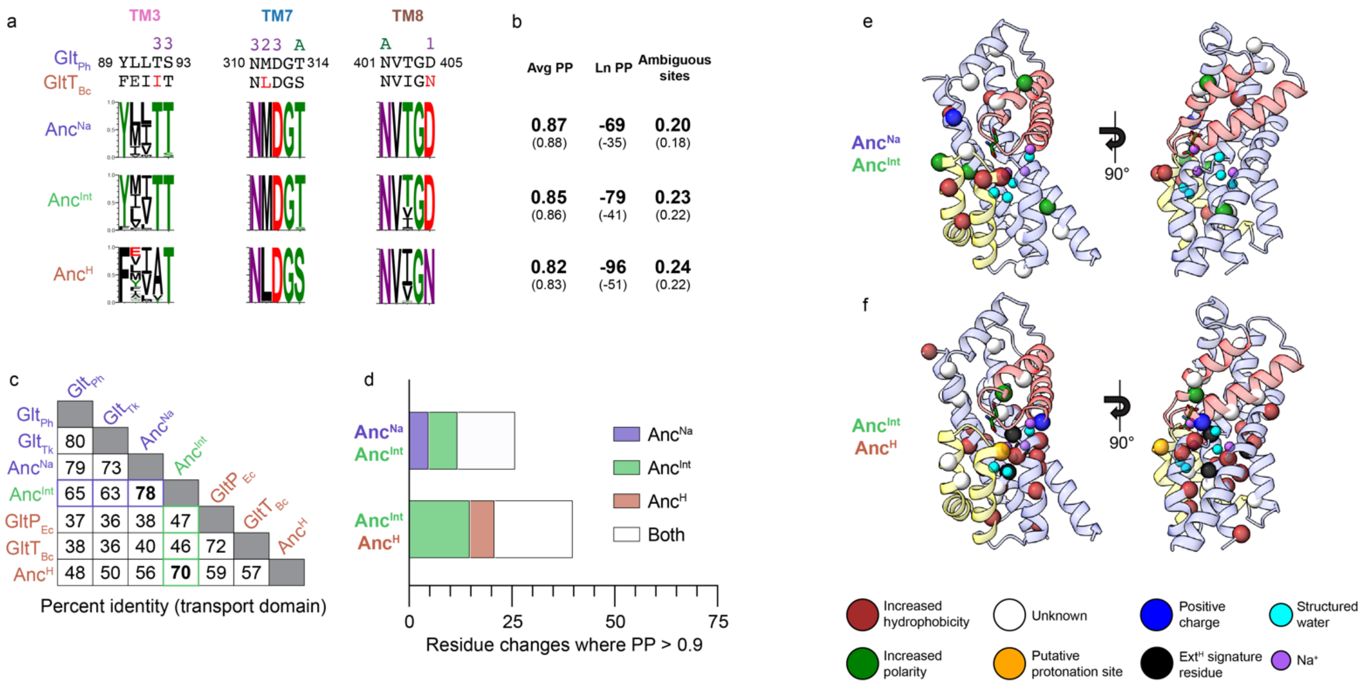

**Supplementary Figure 4. Robustness of the ancestral sequence reconstruction.** (a) Reconstructed ancestral sequences in $Na^+$-binding sites of $Anc^{Na}$, $Anc^{Int}$, and $Anc^{H}$. WebLogos visually represent the posterior probability per site in each ancestral sequence. The topmost residue at each site is the ML residue. $Glt_{Ph}$ and $Glt_{Bc}$ sequence alignments are shown above the WebLogos for reference, along with annotations for L-Asp binding residues (A) or $Na^+$-binding residues (1-3). (b) Average posterior probability (PP) per site, the natural logarithm of the total posterior probability of the entire ancestral sequence (Ln PP), and the fraction of the ambiguous sites (defined as two or more amino acids per site with a probability greater than 0.2) in each ancestor. Bold, larger values are for the entire sequence; values in parentheses are for the transport domain alone. (c) Pairwise identity (in %) matrix of ancestral and extant transporters. The alignment included only transport domains, defined as $Glt_{Ph}$ residues 78-110, 228-416. (d) Visual representation of the changes between two ML ancestral sequences. In each bar, the first segment is the number of changes where the posterior probability is greater than 0.9 in only the first ancestor, i.e., a robustly reconstructed site is ambiguously reconstructed in the second ancestor; the second segment is the number of changes where the posterior probability is greater than 0.9 in only the second ancestor, while the site is ambiguous in the first ancestor; the third segment is the number of changes where the posterior probability is greater than 0.9 in both ancestors with both sites confidently reconstructed. (e-f) Probable sequence changes between $Anc^{Na}$ and $Anc^{Int}$ (e) and $Anc^{Int}$ and $Anc^{H}$ (f), mapped onto the transport domain of $Glt_{Ph}$. Bottom: Labeling scheme for residues. Larger spheres represent sequence transitions; smaller spheres represent $Na^+$ sites and structured waters observed in $Glt_{Ph}$ (PDB 7RCP). “Unknown” indicates when there is no clear physicochemical difference between the two residues. $Ext^{H}$ signature residues are [T/S]92[A/L/I/V/M], M311L, and D405N; putative protonation site is S279E.

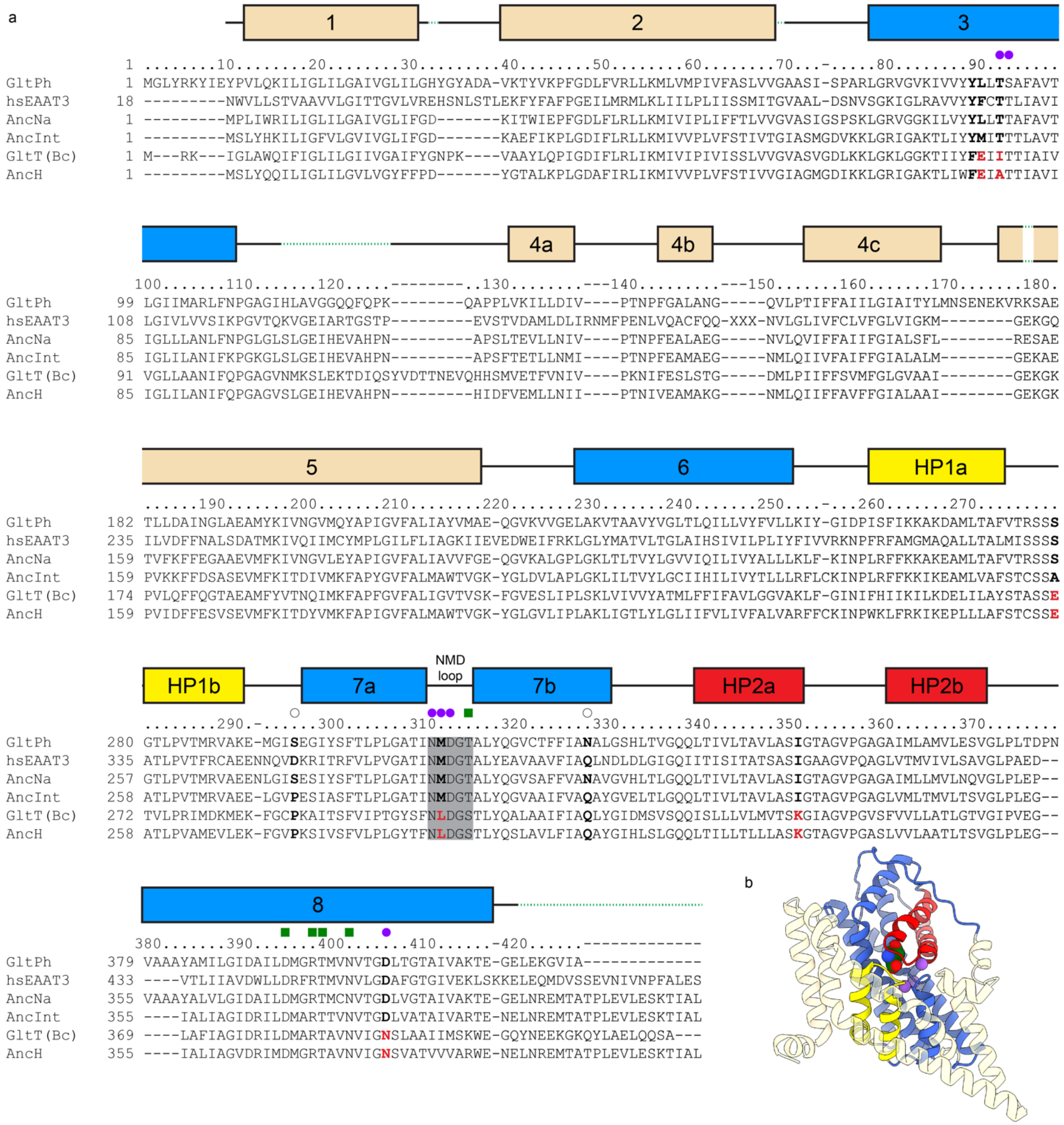


**Supplementary Figure 5. Sequence alignment of extant and ancestral glutamate transporter homologs.** (a) Alignment sequence numbering corresponds to $Glt_{Ph}$ numbering. Extant sequences are $Glt_{Ph}$ (UniProt O59010), $GltT_{Bc}$ (P24944), and human EAAT3 (P43005). TMs, shown above the alignment, are colored wheat for the scaffold/trimerization domain, blue for the transport domain, yellow for re-entrant helical hairpin HP1, and red for re-entrant helical hairpin HP2. Dashed green lines represent residues grafted from an extant sequence into ancestral protein constructs in place of poorly aligned regions (see Methods). Filled purple circles mark residues in $Glt_{Ph}$ that coordinate $Na^+$ ions with their side chains, filled green squares mark residues in $Glt_{Ph}$ that coordinate L-Asp with their side chains, and empty circles represent allosteric mutations that can turn $Na^+$-coupling on and off. Notable sequence changes in $H^+$-coupled transporters compared to $Na^+$-coupled transporters are shown in bold, and red residues are changes found at $Na^+$-binding sites. (b) Structure of a $Glt_{Ph}$ protomer in the outward-facing state (PDB 6UWF), colored as in (a). Purple spheres are $Na^+$ ions (placed based on PDB 6X15), and the substrate is green.

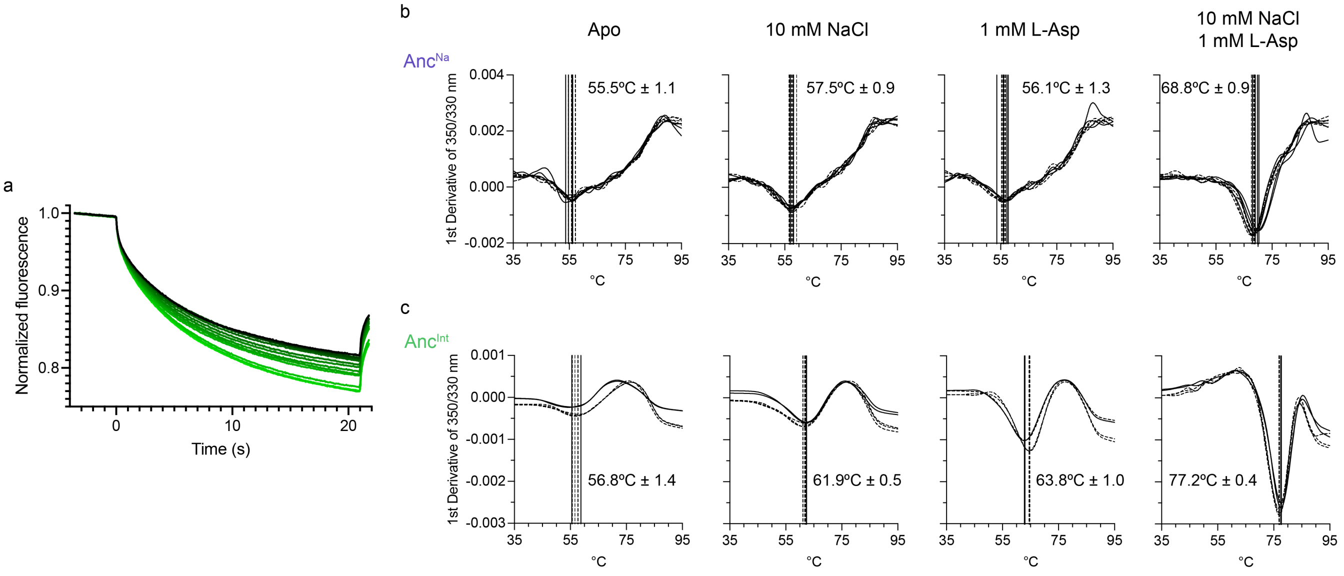


**Supplementary Figure 6. L-Asp and $Na^{+}$ binding to $Anc^{Na}$ and $Anc^{Int}$.** (a) Representative raw MST traces from an individual binding affinity experiment. Traces represent the MST signals of GFP-tagged $Anc^{Int}$ in the presence of 100 mM $Na^{+}$ at various L-Asp concentrations. Lighter green lines correspond to increasing L-Asp concentrations. Fraction-bound values in the main text are calculated between 1.5-5s response time, depending on the individual experiment. (b-c) Raw first-derivative curves to determine thermostability as measured by nanoDSF of $Anc^{Na}$ (b) and $Anc^{Int}$ (c); different pattern lines represent traces performed on independent biological replicates. Vertical lines represent $T_m$-s estimated for individual curves. Mean values with standard deviations are shown on the panels.

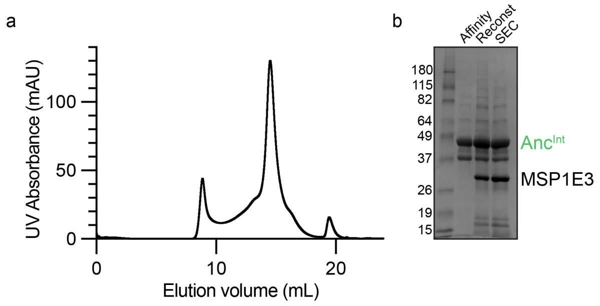


**Supplementary Figure 7. Nanodisc reconstitution of Anc$^{Int}$.** (a) Representative size-exclusion chromatography profile of GFP-free Anc$^{Int}$ reconstituted into MSP1E3 nanodiscs. The column is a Superose 6 Increase 10/300 GL, and the buffer is 20 mM HEPES pH 7.4, 100 mM NMDG-Cl. (b) Representative Coomassie-stained SDS PAGE of GFP-free Anc$^{Int}$ and reconstitution into MSP1E3 nanodiscs. “Affinity” refers to pooled and concentrated elution fractions following Streptactin XT affinity purification. “Reconst” refers to Anc$^{Int}$ successfully reconstituted into MSP1E3 nanodiscs after BioBead application and ultracentrifugation. “SEC” refers to pooled and concentrated peak fractions of Anc$^{Int}$/MSP1E3 eluting at ~14.5 mL following size-exclusion chromatography.

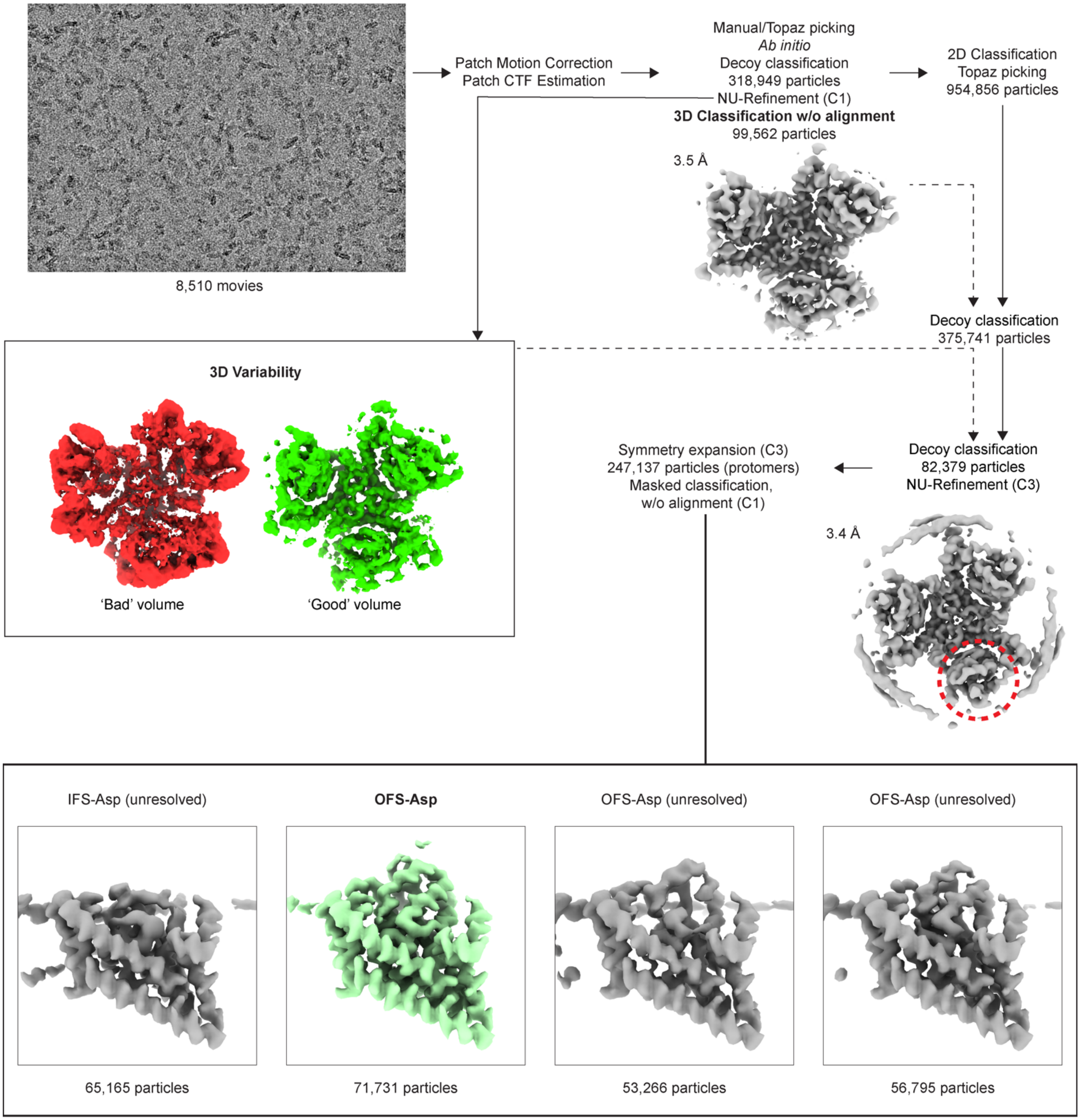


**Supplementary Figure 8. Processing workflow of Anc$^{Int}$, substrate-bound conditions.** Steps are described in further detail in Methods. All steps were performed in cryoSPARC. 'Decoy classification' is a nickname for heterogeneous refinement using 'decoy' noise volumes and a 'good' and 'bad' volumes obtained from 3D Variability (Methods). The dashed red circle is the approximate location of the mask used for masked 3D classification. All maps are unsharpened and contoured to a σ of 10. Colored protomers were used for further local refinement and model building, with the other two protomers removed for clarity.

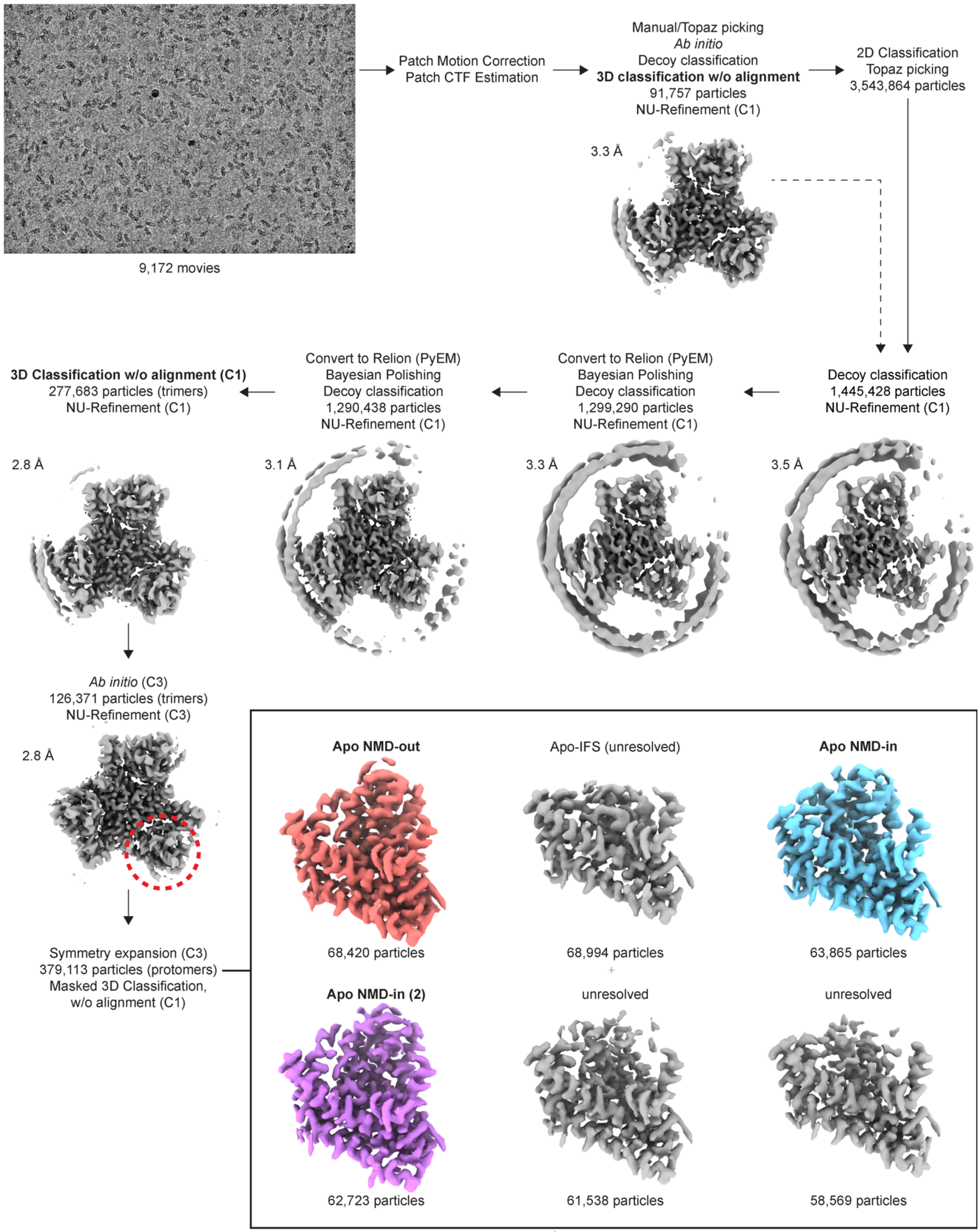


**Supplementary Figure 9. Processing workflow of Anc[Int], apo conditions.** Steps are described in further detail in Methods. All steps were performed in cryoSPARC except for particle stack conversion to Relion (PyEM) and Bayesian Polishing (Relion). 'Decoy classification' is a nickname for heterogeneous refinement using 'decoy' noise volumes (Methods). The dashed red circle is the approximate location of the mask used for masked 3D classification. All maps are unsharpened and contoured to a σ of 10. Colored protomers were used for further local refinement and model building, with the adjacent two protomers removed for clarity.

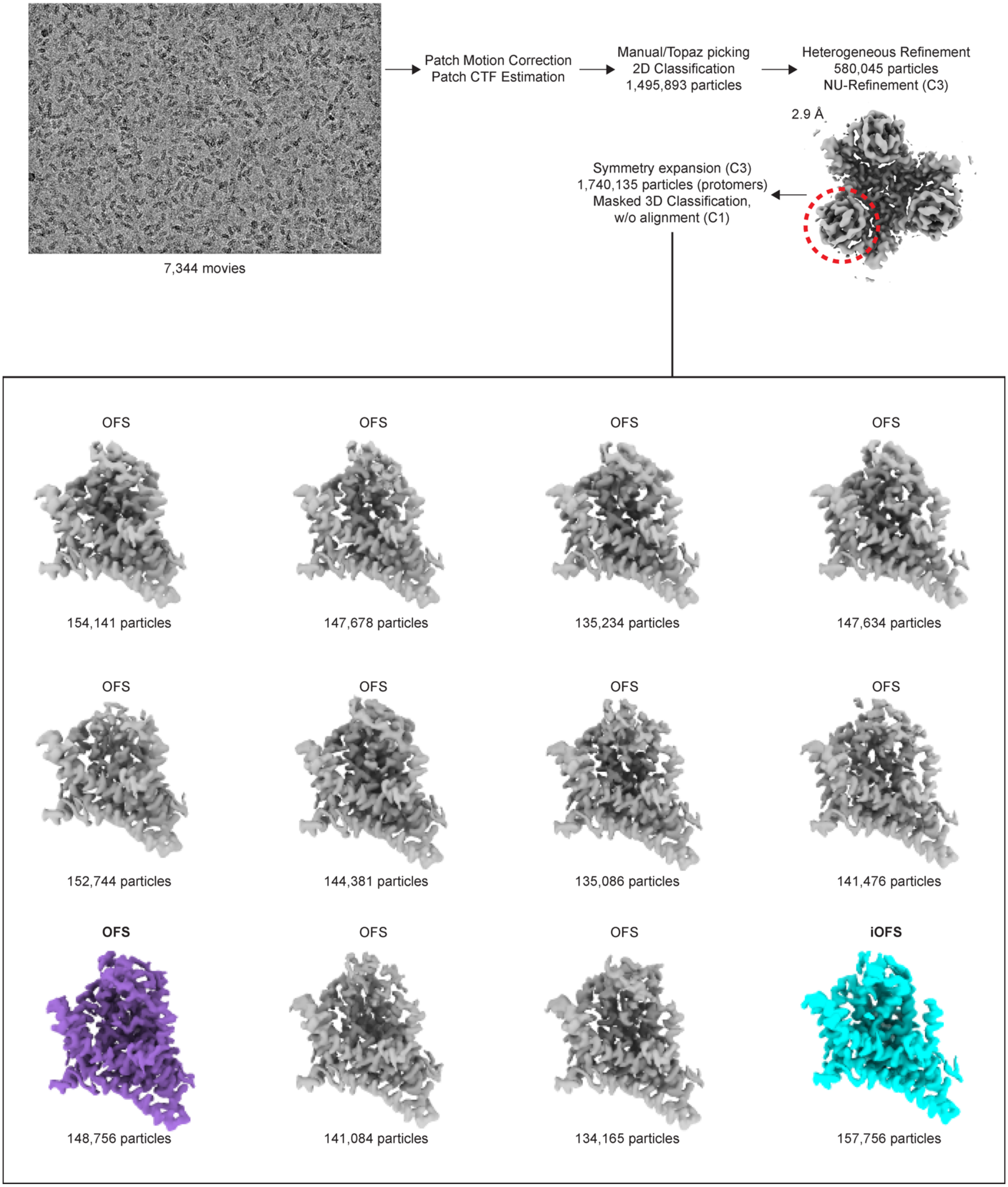


**Supplementary Figure 10. Processing workflow of $Glt_{Ph}$, apo conditions.** Steps are described in further detail in Methods. All steps were performed in cryoSPARC. The dashed red circle is the approximate location of the mask used for masked 3D classification. All maps are unsharpened and contoured to a σ of 10. Colored protomers were used for further local refinement and model building, with the other two protomers removed for clarity.

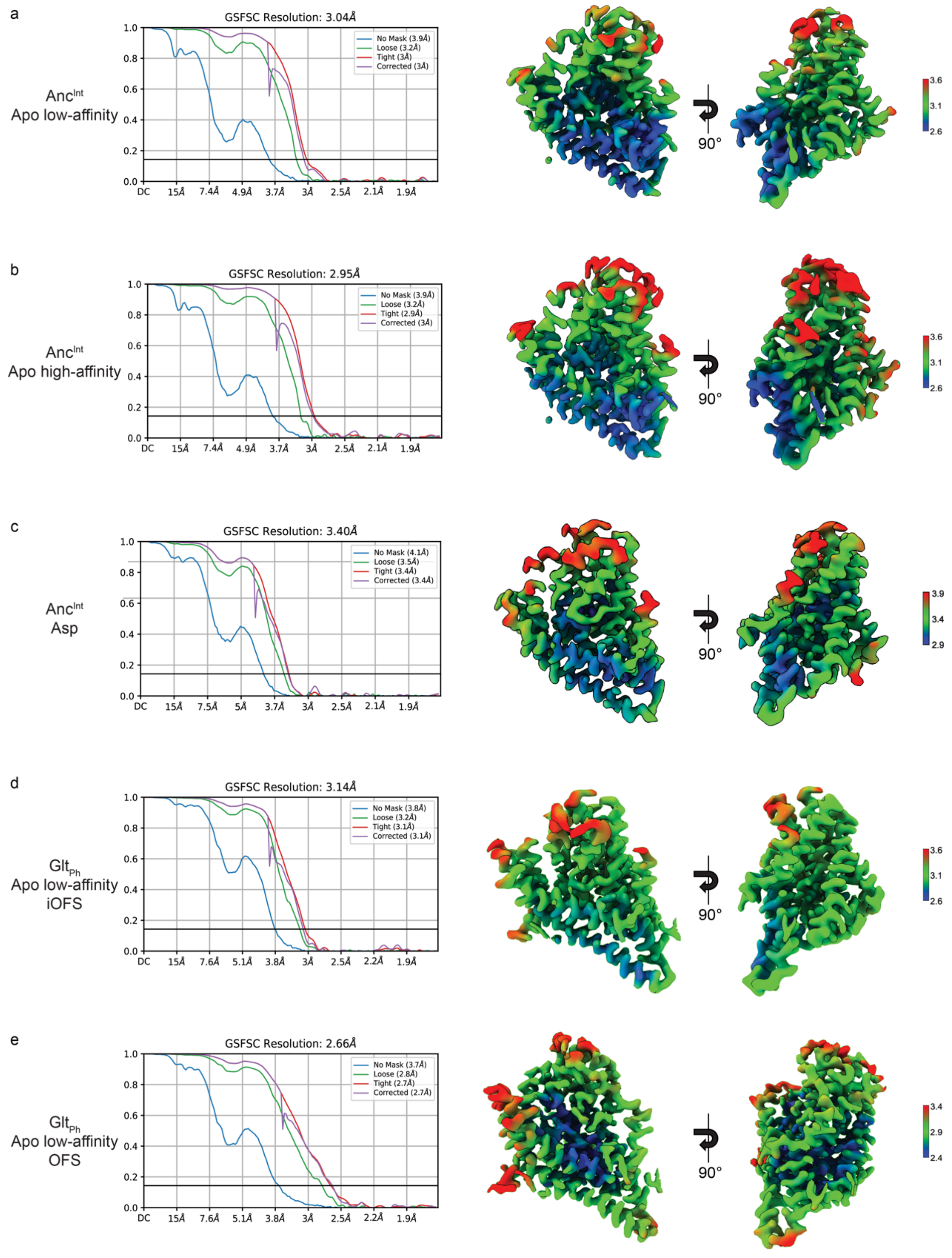


**Supplementary Figure 11. Validation statistics for all structures.** From left to right, map FSC from local refinement in cryoSPARC and local resolution estimation of the unsharpened map. All maps are contoured at 8 σ. The adjacent protomers (unused in refinement) are removed for clarity.

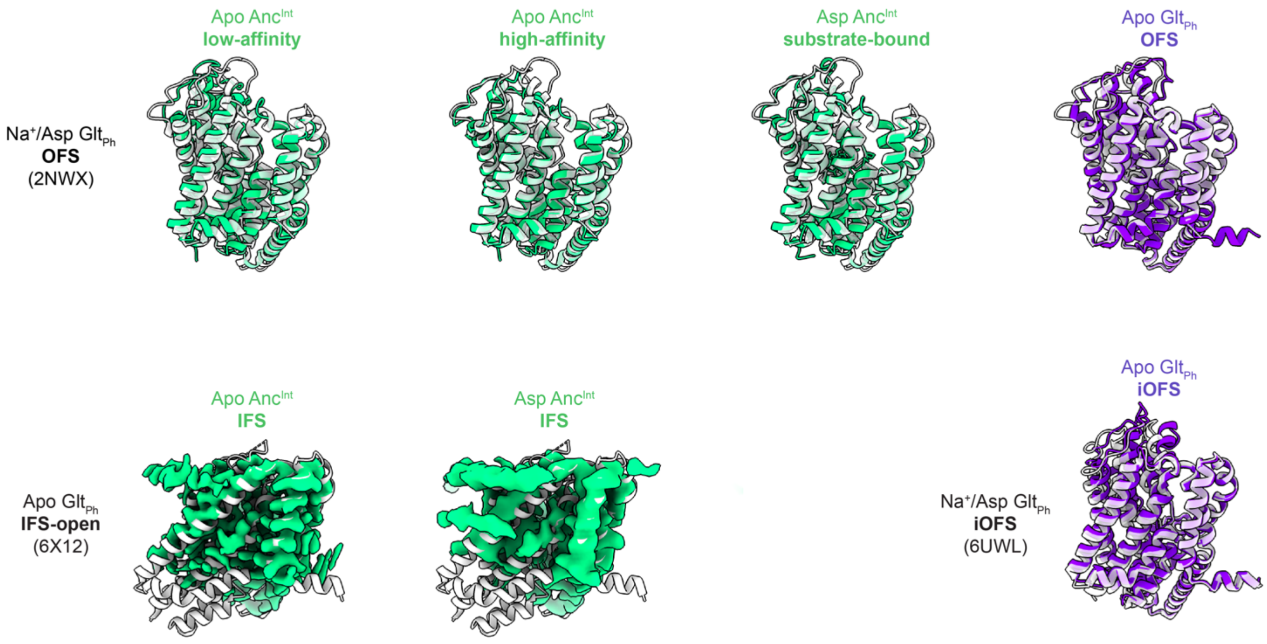


**Supplementary Figure 12. Global transport domain conformations of structures obtained in this study.** $Anc^{Int}$ (green) or $Glt_{Ph}$ (purple) structures were aligned on the scaffold domain (residues 1-75, 140-225) of the closest $Glt_{Ph}$ structure in the PDB (translucent white). These $Glt_{Ph}$ structures were either OFS (2NWX) or iOFS (6UWL). IFS states in $Anc^{Int}$ were incompletely resolved, and models could not be built; therefore, the maps from 3D classification were contoured to a σ of 6 and aligned to the closest $Glt_{Ph}$ structure, IFS-open (white, 6X12).

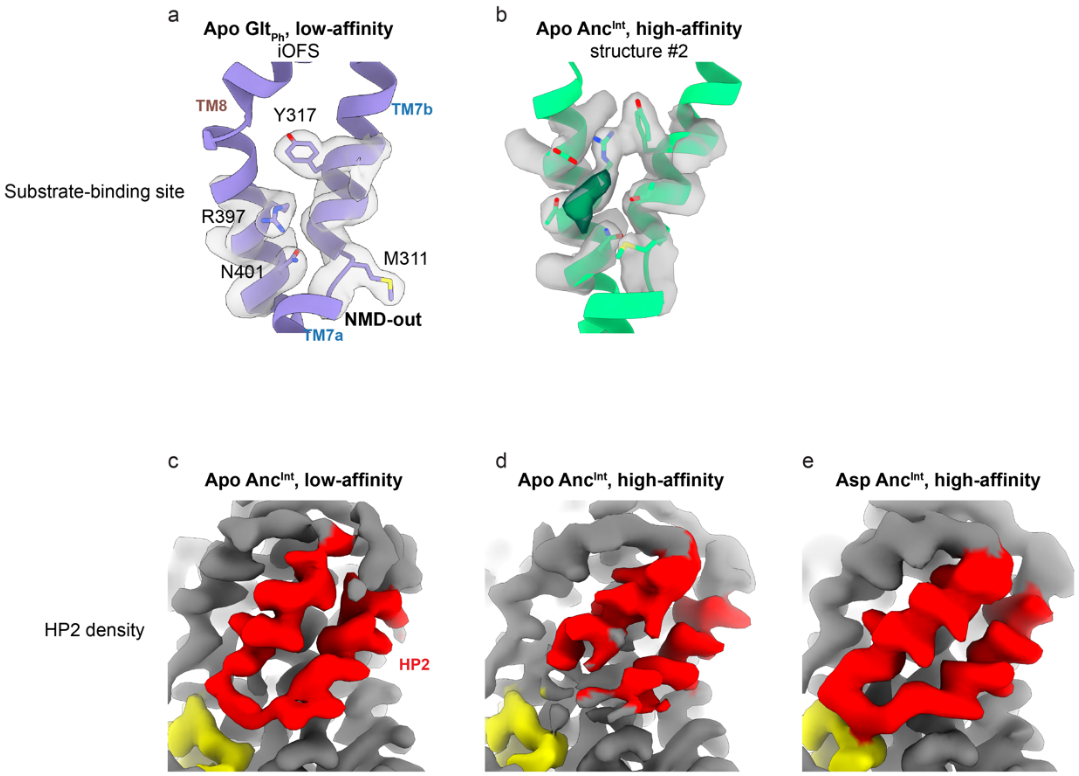


**Supplementary Figure 13. Alternate substrate binding configurations and Anc$^{Int}$ HP2 densities.** (a) Substrate-binding sites of apo-iOFS Glt$_{Ph}$ in the low-affinity configuration. (b) Substrate-binding site of apo Anc$^{Int}$ in the high-affinity configuration #2. The green density overlaps with the L-Asp binding site but is too large and differently shaped to correspond to L-Asp. (c-e) HP2 densities of apo Anc$^{Int}$, low-affinity (c), apo Anc$^{Int}$, high-affinity (d), and L-Asp Anc$^{Int}$, high-affinity (e). Densities are unsharpened and contoured 10 σ. HP1 is in yellow, and HP2 is in red.

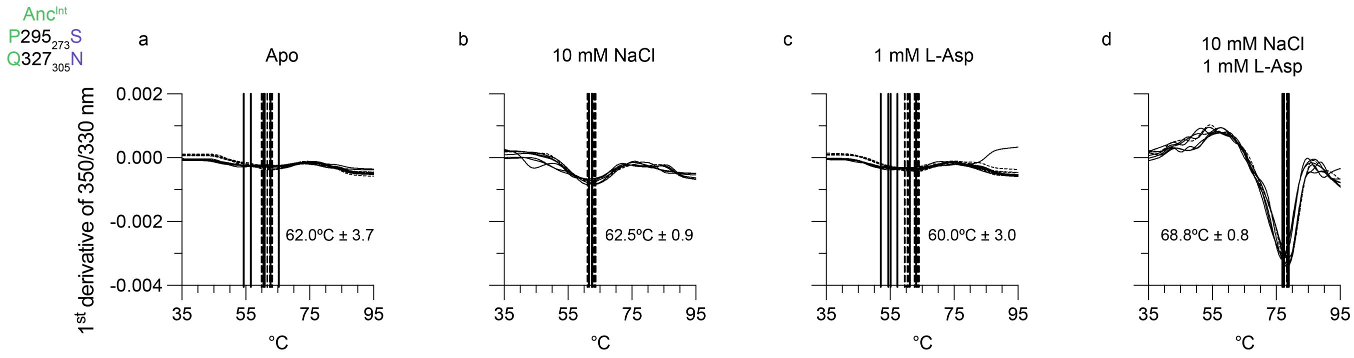


**Supplementary Figure 14. L-Asp and $Na^+$ bind cooperatively to $Anc^{Int}$ $P295_{273}S/Q327_{305}N$.** Raw first-derivative curves to determine thermostability as measured by nanoDSF in apo conditions (a), 10 mM NaCl (b), 1 mM L-aspartate (c), and both NaCl and L-aspartate (d). Different pattern lines represent traces performed on independent biological replicates. Vertical lines represent $T_m$-s estimated for individual curves. Mean values with standard deviations are shown on the panels.

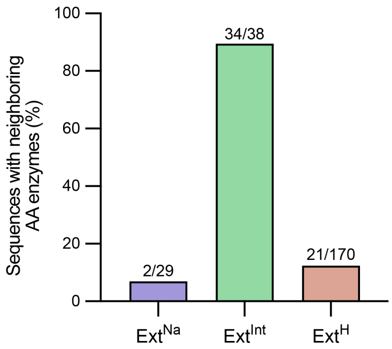


**Supplementary Figure 15. Genomic pairing of uncoupled transporters and enzymes.** Transporters were considered to have a paired enzyme if the enzyme was within 3 protein-coding genes away from the transporter and if the enzyme had putative free amino acid/dicarboxylate modifying function according to Pfam annotation. The fraction of sequences with genomic transporter/enzyme pairs is on the top of each bar. Only transporters with available genomic neighborhood information were considered; for the $Ext^{Na}$, $Ext^{Int}$, and $Ext^{H}$ clades, this corresponds to 29/186, 38/161, and 170/773, respectively.

| **Protein** | **Anc$^{Int}$** | | | **Glt$_{Ph}$** | |
|---|---|---|---|---|---|
| **Conditions** | **apo** | | **L-Asp** | **apo** | |
| *Cryo-EM data acquisition* | | | | | |
| Pixel size (Å/px) | 0.4125 | | 0.4165 | 0.4230 | |
| Frames per movie | 40 | | 50 | 40 | |
| Total dose ($e^-$/Å$^2$) | 50.29 | | 58.00 | 53.42 | |
| Defocus range (μm) | -0.9 to -2.1 | | -0.9 to -2.1 | -0.9 to -2.1 | |
| *Cryo-EM data processing* | | | | | |
| # of movies (after curation) | 9,172 | | 8,510 | 7,344 | |
| Initial # of particles | 3,543,864 | | 954,856 | 1,682,016 | |
| # C3-expanded particles | 379,113 | | 247,137 | 1,740,135 | |
| **Individual structures** | **Low affinity** | **High affinity** | **Asp** | **iOFS** | **OFS** |
| # particles (protomers) | 68,420 | 63,865 | 71,731 | 157,756 | 148,756 |
| Resolution<br>(masked FSC = 0.143, Å) | 3.0 | 3.0 | 3.4 | 3.1 | 2.7 |
| EMDataBank (EMDB) ID | EMD-44526 | EMD-44527 | EMD-44528 | EMD-44529 | EMD-44530 |
| *Model Refinement* | | | | | |
| PDB ID | 9BGY | 9BGZ | 9BH0 | 9BH1 | 9BH2 |
| Model resolution<br>(FSC = 0.5 / 0.143 Å) | 3.2 / 2.9 | 3.2 / 3.0 | 3.6 / 3.3 | 3.4 / 3.1 | 3.3 / 2.7 |
| **Model composition** | | | | | |
| Non-hydrogen atoms | 2,916 | 2,783 | 2,935 | 2,942 | 3,046 |
| Protein residues | 389 | 369 | 391 | 395 | 414 |
| Ligands | 0 | 0 | 1 | 0 | 0 |
| **R.m.s. deviations** | | | | | |
| Bond lengths (Å) | 0.002 | 0.003 | 0.002 | 0.003 | 0.003 |
| Bond angles (°) | 0.499 | 0.542 | 0.573 | 0.482 | 0.523 |
| **Validation** | | | | | |
| MolProbity score | 1.36 | 1.42 | 1.26 | 1.30 | 1.37 |
| Clash score | 6.28 | 7.60 | 4.94 | 5.60 | 6.79 |
| Poor rotamers (%) | 0.32 | 0.66 | 0.32 | 0.00 | 0.62 |
| **Ramachandran plot** | | | | | |
| Favored (%) | 97.93 | 98.35 | 98.20 | 99.74 | 98.30 |
| Allowed (%) | 2.07 | 1.65 | 1.80 | 0.26 | 1.70 |
| Disallowed (%) | 0.00 | 0.00 | 0.00 | 0.00 | 0.00 |

**Supplementary Table 1.** Cryo-EM acquisition parameters, data processing metrics, and model refinement/ validation statistics.
